# Pro-myelinating Clemastine administration improves recording performance of chronically implanted microelectrodes and nearby neuronal health

**DOI:** 10.1101/2023.01.31.526463

**Authors:** Keying Chen, Franca Cambi, Takashi D.Y. Kozai

## Abstract

Intracortical microelectrodes have become a useful tool in neuroprosthetic applications in the clinic and to understand neurological disorders in basic neurosciences. Many of these brain-machine interface technology applications require successful long-term implantation with high stability and sensitivity. However, the intrinsic tissue reaction caused by implantation remains a major failure mechanism causing loss of recorded signal quality over time. Oligodendrocytes remain an underappreciated intervention target to improve chronic recording performance. These cells can accelerate action potential propagation and provides direct metabolic support for neuronal health and functionality. However, implantation injury causes oligodendrocyte degeneration and leads to progressive demyelination in surrounding brain tissue. Previous work highlighted that healthy oligodendrocytes are necessary for greater electrophysiological recording performance and the prevention of neuronal silencing around implanted microelectrodes over chronic implantation. Thus, we hypothesize that enhancing oligodendrocyte activity with a pharmaceutical drug, Clemastine, will prevent the chronic decline of microelectrode recording performance. Electrophysiological evaluation showed that the promyelination Clemastine treatment significantly elevated the signal detectability and quality, rescued the loss of multi-unit activity, and increased functional interlaminar connectivity over 16-weeks of implantation. Additionally, post-mortem immunohistochemistry showed that increased oligodendrocyte density and myelination coincided with increased survival of both excitatory and inhibitory neurons near the implant. Overall, we showed a positive relationship between enhanced oligodendrocyte activity and neuronal health and functionality near the chronically implanted microelectrode. This study shows that therapeutic strategy that enhance oligodendrocyte activity is effective for integrating the functional device interface with brain tissue over chronic implantation period.

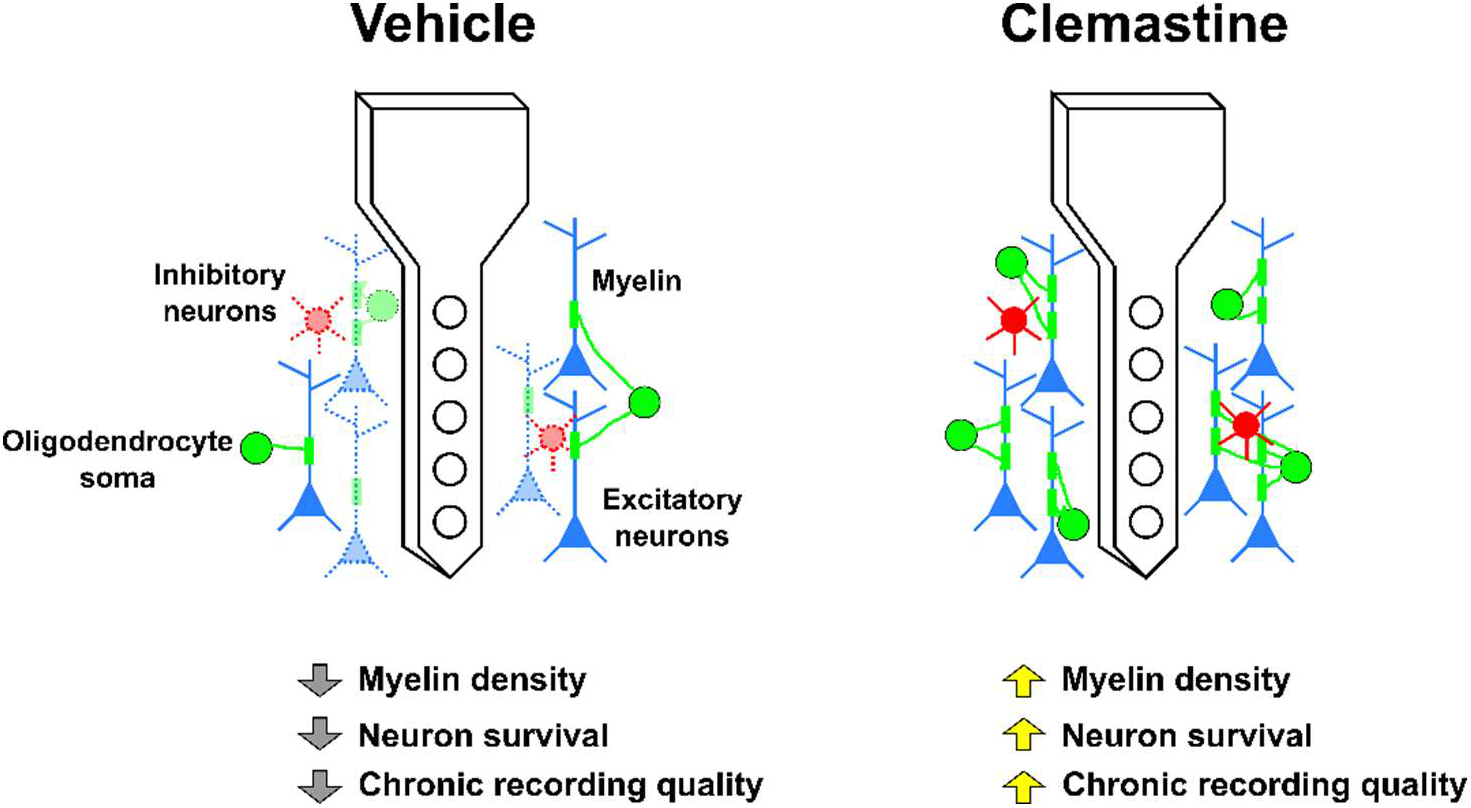

## 1. Introduction

Penetrating intracortical microelectrodes that monitor and/or modulate surrounding neural tissue are front-end components of brain-machine interfaces used to investigate basic neuroscience and clinically treat neurological disorders [1–7]. However, the biological responses to long-term microelectrode implantation, including blood-brain barrier rupture, microglia activation, and astrocyte reactivity, can lead to local neurodegeneration and scar tissue formation resulting in gradual decline of the device’s functional performance [8–15]. The variability in the loss of signal stability and sensitivity over the chronic implantation period further limits the applications of these intracortical microelectrodes [9, 16–19]. Therefore, investigations to improve the long-term performance of these microelectrodes have focused on reducing the tissue response caused by the chronic implantation of microelectrodes [20–27].

Many studies attempted to improve performance by mitigating microglia activation and astrocyte reactivity [20, 24, 28–33]. Some treatments that reduce glial activation have enhanced recording quality, such as increased detection of single-unit action potentials and improved signal strength [34]. However, even when there is robust neural density around the implant and minimal microglial and astrocyte encapsulation, microelectrodes can still fail to record action potentials due to silencing of nearby neurons [35, 36]. These observations highlight that recording performance cannot be exclusively explained by microglia, astrocytes, and neurons and suggests that additional cellular players could be key drivers for microelectrode functional performance. Oligodendrocytes compose 45-75% of glial cells [37, 38] but have been largely understudied in the neural engineering field.

Oligodendrocytes differentiate from oligodendrocyte precursor cells (OPC) and generate myelin sheaths around axons [39–41], leading to large surface area contacts with these neurons. Through these intimate contacts, oligodendrocytes regulate neuronal activity and function by enhancing action potential propagation, and, importantly providing metabolic and neurotrophic support [42–46]. Specifically, oligodendrocytes facilitate saltatory conduction via insulating properties of the lipid-rich myelin processes [47–49]. Furthermore, oligodendrocytes regulate metabolic coupling between axons and myelin, discussed in detail in [42, 44]. Briefly, oligodendrocytes provide metabolites to neurons when under heavy metabolic burden, since neurons have no glycogen storage [42, 44, 50, 51]. Interestingly, while the oligodendrocyte activity is tightly influenced by neural network activity[52–54], enhanced oligodendrocyte activity leads to improvement of brain network functionality. For example, pharmacologically enhanced myelination has been shown to improve memory recall and hippocampal network activation [55–57]. Therefore, maintaining oligodendrocyte activity is critical for neuronal health and functionality as well as circuit connectivity over a large distance.

However, oligodendrocytes are vulnerable to traumatic brain injuries and neuroinflammation [47, 58–67]. High levels of pro-inflammatory cytokines and chemokines, oxidative stress, metabolic stress, or excitotoxic glutamate release are associated with oligodendrocyte degeneration, demyelination, and OPC differentiation failure in multiple brain injury models and neurodegenerative diseases, such as multiple sclerosis [59, 61, 63, 64, 67–72]. Deficits in oligodendrocyte lineage cells lead to axonal pathology, impairment of network activation, and ultimately behavioral dysfunction [49, 56, 70, 73, 74]. Microelectrode implantation creates a similar local inflammatory injury site that causes oligodendrocyte loss, demyelination, and ineffective OPC turnover in the microenvironment near the implant [59]. Depleting oligodendrocytes with the toxin cuprizone led to immediate decline of recording performance that persisted throughout the entire recording period [75]. Cuprizone-induced oligodendrocyte depletion did not affect the neuronal density around the implants, suggesting neuronal silencing occurs in the microenvironment around the microelectrode. Given the multiple mechanisms of neuronal support mediated by OL, their loss would affect neurons in multiple ways: 1) reduced metabolic support [76], 2) loss of neurotrophic growth factors [77], 3) reduced conduction velocity [78], 4) faster action potential degradation across axons [42, 51]. Therefore, targeting an individual mechanism would likely not be sufficient to rescue OL-mediated neuron support. In these studies, we sought to investigate whether enhancing oligodendrocyte cell density and function would enhance functional recording performance over the chronic implantation period.

One potential pharmacological candidate that targets oligodendrocytes and promotes myelination is Clemastine [56, 57, 62, 79–83]. This drug is a Food and Drug Administration (FDA)-approved over the counter antihistamine which also has pro-myelinating effects [83]. Clemastine selectively enhances OPC differentiation into mature oligodendrocytes and integrates new myelin into the functional network, via binding to the M1 muscarinic receptors on OPCs [62]. While many studies have shown Clemastine’s beneficial effects on neurodegeneration models improving learning, working memory, social disorders, no significant changes in morphology and density of axons or neurons have been observed [54–56, 62], suggesting that Clemastine’s profound therapeutic impact on neural network activity is indirectly mediated by oligodendrocyte lineage cells. Meanwhile, Clemastine has low side effect profile with rarely reported adverse events [82]. Therefore, we selected Clemastine to enhance oligodendrocytes and assess its effect on chronically implanted microelectrode functional performance.

In this study, we tested the hypothesis that Clemastine administration leads to a higher microelectrode chronic recording quality in the cortex and hippocampal CA1 relative to vehicle treated controls over a 16-week implantation period. We evaluated Clemastine’s impact on preserving network functionality in the microenvironment around the implant compared to vehicle condition, including the neuronal functional subtype recording viability, firing properties, and LFP synchronization between different laminar structures. We found that Clemastine significantly increased oligodendrocyte density and myelination at the end of the 16-week study. Microelectrode recording performance over the implantation period was significantly improved compared to the vehicle controls. Detailed electrophysiological analyses revealed that Clemastine improved the functional activity of different neuronal subtypes in a depth-dependent manner and resulted in a significantly stronger laminar connectivity especially over 13-16 weeks post-implantation. Overall, we demonstrate that targeting oligodendrocyte lineage health as a novel therapeutic strategy is effective in improving neuronal recording viability and functionality, which in turn improves chronic functional recording performance.

## 2. Methods

C57BL/6J mice receiving Clemastine administration and vehicle solutions (each condition N =8, 4 males and 4 females) were implanted with the microelectrodes over 16 weeks. All animal care and procedures were performed under approval of the University of Pittsburgh Institutional Animal Care and Use Committee and in accordance with regulations specified by the Division of Laboratory Animal Resources.

### 2.1 Clemastine administration

Adult 6-8 week old C57BL/6J mice (Jackson Laboratory, Bar Harbor, ME) were given either Clemastine (dissolved in 10% DMSO/PBS, 10mg/Kg body weight; Tocris Bioscience) or vehicle solution (only 10% DMSO/PBS) daily through intraperitoneal injection. The control group is always selected on an experiment-by-experiment basis based on the specific scientifically guided question rather than maintaining uniformity across programs, which would introduce mixed variable effects that limits the ability to interrogate and interpret the effect of the treatment. Here, the drug, Clemastine, was administered via intraperitoneal injection in a DMSO/PBS vehicle solution delivered via an injection instead of it being mixed in with the food chow, like Cuprizone in a previous study [75]. Therefore, the appropriate control group to control for the effect of the drug delivery mechanism in this study is a DMSO/PBS vehicle intraperitoneal injection without the Clemastine drug. Using a control “chow” vehicle would not control for the stress that the Clemastine group might experience during intraperitoneal injection. Ensuring that both experimental and control group experience the same injection conditions for these variables allow for a fairer evaluation of the effects of Clemastine alone instead of the effects of Clemastine and intraperitoneal injection related effects. To ensure that Clemastine has effect during acute recording sessions, we start administration 7 days prior to implant according to previous reports [62, 79, 83]. Evidence has shown that consecutive 7 days Clemastine administration can effectively rescue behavioral deficits from neurological disorder models [62, 79, 83]. The pretreatment of Clemastine primes the oligodendrocyte lineage cells for the implantation injury, which occurs at a previously predetermined scheduled time (unlike unscheduled traumatic brain injuries). Mice received treatment of either Clemastine (in the vehicle) or vehicle (without Clemastine) for 7 days prior to the microelectrode implantation surgery and then daily treatments were maintained for another 16 weeks for the recording experiments. All experimental mice were housed in a temperature-controlled, humidity-controlled, and 12 h light/dark cycle facility. Food and water were available ad libitum.

### 2.2 Microelectrode implantation surgery

The implantation surgery procedure has been previously described [58, 84]. Mice were anesthetized with a cocktail of xylazine (7 mg/kg) and ketamine (75 mg/kg). After fixing the animal onto a stereotaxic frame, a 1-mm-square craniotomy was performed over the left visual cortex, with the center 1.7 mm lateral to midline and 2.3 mm posterior to bregma. Michigan-style functional single-shank microelectrode arrays (A16-3 mm-100-703-CM15, NeuroNexus, Ann Arbor, MI) were perpendicularly inserted into the craniotomy while avoiding surface vasculature [85]. The ground wire was wrapped around a bone screw over ipsilateral motor cortex and the reference wire was wrapped around a bone screw over contralateral visual cortex. During the surgery, the body temperature and respiration were monitored. Post-operative ketofen (5 mg/kg) were given on the surgery day and following two consecutive days.

### 2.3 Electrophysiological recording procedure & Electrochemical impedance spectroscopy (EIS)

Electrophysiological recording and EIS were performed inside a grounded Faraday cage to avoid environmental noises. Awake mice were head-fixed on a spinning disk platform with the microelectrode headstage connected. The electrophysiological data was sampled at 24,414 Hz (RZ2, Tucker-Davis Technologies, Alachua, FL) for spontaneous activity and was performed in a blackout chamber. The visual stimulation data was collected with the contralateral eye positioned 60° relative to the monitor to maximize visual stimulation to the mouse’s visual field. Sixty-four drifting grating visual stimulation were presented as 1-s on and 1-s off in 8-directions, programmed in MATLAB using the Psychophysics toolbox as described before [58, 84].

For EIS measurements, the headstage of awake mice was connected to an Autolab potentiostat with a 16 channel multiplexer (PGSTAT 128N, Metrohm, Netherlands). Impedances were recorded for each channel using a 10 mV RMS sine wave in a range of 10 Hz to 32 kHz. Here, 1 kHz impedance averaged over all animals was reported for each day.

### 2.4 Recording data analysis

Cortical layer alignment of the microelectrode array for each animal was performed for every recording session using current source density (CSD) analysis and centered onto Layer 4 [84]. A 2^nd^ order Butterworth filter at 0.4-300 Hz was applied to obtain the local field potential (LFP) data stream. LFP signals were smoothed by running 1-dimensional line fit. Then CSD was constructed by computing the second spatial derivative of evoked (stimulus-locked) LFP for each electrode site 100 μm spacing along the microelectrode shank [84]. Then CSD was averaged across 64 trials of visual stimulation and the location of the first current sink (minimum value) within 100 ms was determined as L4 depth. All depth-dependent recording metrics was normalized to the corresponding L4.

#### 2.4.1 Single-unit sorting and analysis

Broadband electrophysiological raw data was processed using a custom MATLAB script described previously [86]. A 2^nd^ order bandpass Butterworth Filter was applied to produce 0.3-5 kHz spike stream. Common average referencing (CAR) was applied to normalize the data streams [87]. Candidate single-units (SUs) were detected by a negative threshold of 3.5 standard deviation from the mean and further discriminated by principal component analysis (PCA). SUs were then manually sorted by evaluating the waveform shapes, auto-correlograms, and peri-stimulus time histograms (PSTH) with 50-ms bins. ‘SU yield’ was the percentage of channels that contain at least one SU. ‘Single-to-noise ratio (SNR)’ of each SU was defined as the ratio of peak-to-peak amplitude of the waveform over the noise floor. ‘Noise floor’ for each channel was calculated as 2 standard deviations of the spike data stream after removing all thresholding events. ‘Averaged SNR’ was calculated as the mean of the largest SNR of each channel across the microelectrode array, where SNR = 0 when a SU was not detected. The ‘average SNR/active site’ was the mean SNR only for channels that detected a SU.

#### 2.4.2 Multi-unit analysis

As previously described [58, 84], the multi-unit activity (MUA) was defined as the activity of all threshold-crossing events, including both SU and outlier clusters. The firing rate of MU activity was calculated as the average number of MU events during the 1-s visual stimulation period or pseudotriggers over an equivalent period of time during spontaneous (or resting state) condition with the monitor turned off. To evaluate the temporal responsiveness of MUA to visual stimulus within 1s, the ‘multi-unit (MU) yield’ and ‘signal-to-noise firing rate ratio (SNFRR)’ were compared between prestimulus OFF period and 1-s visual stimulation ON period. MU yield was calculated as the percent of electrode sites that have a significant (p < 0.05) different MUA firing rate during stimulation ON compared to OFF (right before stimulus onset). Similarly, the SNFRR quantitatively described the increase in firing rate of MU activity in response to visual stimuli, which was calculated as the difference in the MU firing rates during and after the stimulation normalized to the average standard deviation between each stimulus condition.

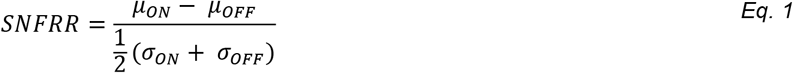

where *μ_ON_* and *μ_OFF_* are the average firing rates (across 64 trials) during visual stimulus ON and OFF conditions, while *σ_ON_* and *σ_OFF_* are the standard deviations of firing rates during ON and OFF conditions.

MU yield and SNFRR require comparisons of MUA before and after the stimulus onset. Since the MUA counts analysis depends on the varying temporal bin size (B) and latency after stimulus onset (L) from 0 to 1 s in length via 1-ms increments, the MU yield and SNFRR were first calculated for all combinations of B and L with prerequisite that;

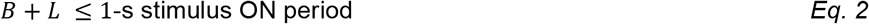

Then B was fixed to the value that optimized the MU yield averaged across all time points in vehicle animals. The differences in temporal responsiveness of visual evoked MUA between Clemastine and vehicle controls was quantified by the latency from the stimulus onset, when bin sizes were fixed in cortex and hippocampus CA1, respectively.

#### 2.4.3 Local field potential power analysis

A 2^nd^ order Butterworth filter at 0.4-300 Hz was applied to obtain the local field potential (LFP) data stream. A multitaper method of 1-s duration, 1-Hz bandwidth, and a taper number of 1 was utilized to produce LFP power spectra. Power spectrum during evoked session was normalized to spontaneous power spectrum as

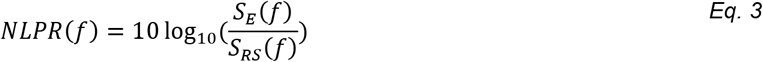

where *NLPR*(*f*) is the normalized LFP power response ratio. S_E_(f) is the visually evoked power spectra, and S_RS_ is the spontaneous power spectra.

#### 2.4.4 Putative excitatory or inhibitory subtype classification of single-units

The sorted SUs were classified based on the action potential waveform shapes, which has been described in [88, 89]. The SU waveform width was defined as the trough-to-peak latency (TP latency), which was the duration between the valley and the peak of the sorted SU waveforms. Following the bimodal distribution of SUs based on the TP latency, the single-units with TP latency > 0.41 ms were tentatively classified as putative excitatory neurons and those with TP latency ≤ 0.41 ms were classified as putative inhibitory neurons.

#### 2.4.5 Laminar coherence analysis

The intra- and interlaminar activity was evaluated by coherence, which quantitatively described the similarity between two LFP signals in the frequency domain [90]. Coherence calculations were performed during the 1-s stimulation period or 1-s spontaneous pseudotrigger, at a half-bandwidth of 3 Hz and a taper number of 5, and then averaged across all trials. Coherence was calculated as

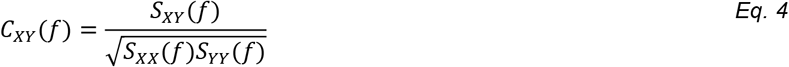

where, *C_XY_*(*f*) is the coherence, *S_XY_*(*f*) is the cross-spectrum of LFP activity between two different channels *X* and *Y*, *S_XX_*(*f*) and *S_YY_*(*f*) are the autospectragram for individual channels.

The changes in circuit connectivity across different depths between the visual stimulation and the prestimulus OFF period was determined by the delta conference Δ*C_XY_*(*f*),

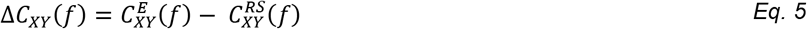

where 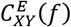 is the visually evoked coherence, and 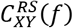 is the spontaneous coherence.

#### 2.4.6 Phase amplitude coupling analysis

Circuit connectivity can be directionally quantified by LFP oscillatory synchronization across difference brain depth, which is the phase-amplitude coupling (PAC) measurement. As a cross-frequency coupling measurement, PAC describes the degree of LFP synchronization connectivity for how well the phase of low frequency oscillations drive the amplitude of coupled high frequency oscillations. The PAC modulation index (MI), which indicate the level of LFP cross frequency synchronization, was calculated based on the Kullback–Leibler (KL) distance formula [91]. The details of MI calculations were described in [91]. Briefly, raw signal was filtered to specific LFP frequencies, *f* (see below for ranges). Then, a Hilbert transform was applied to extract the time series of the phase component as well as the amplitude component from the LFP activity. The phases of slow LFP in channel X was denoted as *Φ_x_* (*t,f_X_*), where *t* represented the time within 1 s following stimulation onset, and *f_X_* represented the low frequency oscillation (*f_X_* = 4, 4.5, 5… 7.5 Hz). The amplitude envelop of high frequency LFP oscillation in another channel Y was denoted as *A_Y_* (*t,f_Y_*), where *f_Y_* represented the high frequency oscillation (*f_Y_* = 30, 30.5, 31… 90 Hz).

Then the composite time series (*Φ_X_* (*t,f_X_*), *A_Y_* (*t,f_Y_*)) was constructed to give the amplitude of LFP oscillation in channel Y at each phase of channel X LFP oscillation. Next, the phases *Φ_x_* (*t,f_X_*) were binned every 18°(20 bins in total). The average of amplitude *A_Y_* (*t,f_Y_*) over each phase bin (*i*) was calculated as *A_Y_* (*t, f_Y_)_φ_X_(t,f_X_)_* (*i*). Finally, the average amplitude *A_Y_* (*t,f_Y_*)*_φ_X_(t,f_X_)_* was normalized by the sum over all bins;

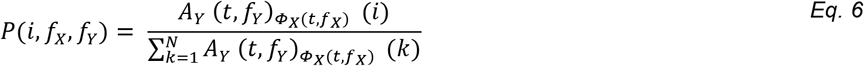

where *P*(*i, f_X_, f_Y_*) is the normalized amplitude distribution over phases, and *N* is the number of phase bins. if there is no PAC between two channels *X* and *Y*, the normalized amplitude distribution *P*(*i, f_X_, f_Y_*) over phase bins would be uniform. The existence of PAC was quantified by the level of deviation of the amplitude distribution *P*(*i, f_X_, f_Y_*) from the uniform distribution, which use KL distance formula to calculate the PAC modulation index (MI). KL distance formula is related to joint entropy (*H* (*f_X_, f_Y_*)), which was calculated as

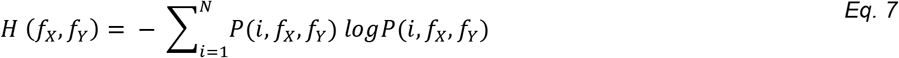

If the normalized amplitude distribution *P*(*i,f_X_,f_Y_*) was uniform, the joint entropy reaches its maximum as *H_o_* = *logN*^2^. Finally, the KL distance was calculated as the difference between *H* (*f_X_*, *f_Y_* *H_o_*, and the MI was defined as the value by dividing the KL distance of the *P*(*i,f_X_,f_Y_*) from the uniform distribution *H_o_*.

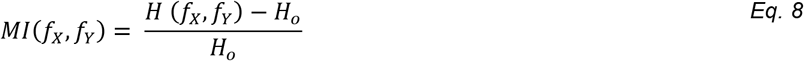

MI was reported as a value between 0 and 1, with a larger MI value indicating a stronger coupling between Channel X low frequency phase and Channel Y high frequency amplitude.

### 2.5 Post-mortem histological analysis+

At the end of 16 weeks implantation, mice were administered a mixture of xylazine (7 mg/kg) and ketamine (75 mg/kg). Deeply anesthetized mice were perfused transcardially with ~100ml 1x PBS to flush the circulating blood and following ~ 50 ml 4% paraformaldehyde (PFA) until tissue fixation was observed. Brains were extracted and postfixed with 4% PFA overnight at 4°C, and then rehydrated in 30% sucrose. Afterward, brains were embedded with the optimum cutting temperature (OCT) media and sectioned horizontally at 25 μm from the cortex surface. Slides were stored at −20°C.

#### 2.5.1 Immunohistochemistry

Standard immunohistochemical staining techniques were performed [58]. Heat-induced antigen retrieval was first performed by sodium citrate buffer (0.1 M citric Acid, 0.1 M sodium citrate). The endogenous peroxidase activity was blocked by following 30% hydrogen peroxide (20 mins). Subsequent blocking was performed using 0.1% Triton-X with 10% normal goat serum in PBS at room temperature for 1 h, to increase the permeability of the brain tissue. Then incubation of primary antibodies to MBP (1:500, Abcam, ab7349), NG2 (1:500, Sigma Aldrich, AB5320), CC1 (1:100, Millipore, OP80), MOG (1:100, Fisher, AF2439), GAD67 (1:500, Abcam, ab213508), CamKiiα (1:100, Abcam, ab22609), MAP2 (1:1000, Abcam, ab5392), NF200 (1:250, Sigma Aldrich, N5389), MCT1 (1:100, Abcam, ab93048), APP (1:400, Abcam, ab2084) overnight at 4°C. After washing with PBS, sections were then incubated with the secondary antibodies (donkey anti-rat 405, Abcam, ab175670, donkey anti-rabbit 488, Abcam, ab150061, donkey anti-mouse 568, Abcam, ab175700, donkey anti-goat 647, Abcam, ab150135, donkey anti-chicken 647, Jackson ImmunoResearch, 730-605-155, Nissl 435/455, Thermo Fisher, N-21479) diluted at 1:500 for 2h at room temperature. Sections were washed and mounted with Fluoromount-G media (SouthernBiotech, #0100-20).

#### 2.5.2 Confocal imaging and data analysis

Confocal microscope (FluoView 1000, Olympus, Inc., Tokyo, Japan) with 20x oil-immersive objective lens was used to capture the TIFF images of probe site and an equivalent area in contralateral sides. The images were carefully acquired in resolution of 16-bit (635.9 × 635.9 μm, 1024 × 1024 pixels) with HiLo setting assistance. A previously published MATLAB script, I.N.T.E.N.S.I.T.Y. was applied to evaluate the intensity of fluorescent markers (MBP/NG2/MOG/MAP2/NF-200/MCT1/APP) binning away from the probe site [92, 93]. Once the probe hole was identified, bins spaced 10 μm apart up to 300 μm away from the probe were generated. The average grayscale intensity was calculated as the mean value of all pixels above the threshold of 1.5 standard deviations above the background noise. The cell counting analysis was performed for CC1/GAD67/CamKiiα. The bin size was modified to 50 μm steps and measured up to 300 μm away from the probe. The cell density was calculated as the total cell counts divided by the tissue area per bin after excluding lost tissue in each bin. For counting neuronal subtypes, GAD67 or CamKiiα were merged with neuronal nuclei marker Nissl and quantified as a functional of distances. The immunohistochemical data was averaged across the animals and plotted as a function of distance away from the probe. The analysis in the contralateral side was measured using the center of the image and averaged over all distance bins.

### 2.6 Statistics

Significant differences between Clemastine and vehicle conditions in recording metrics were determined by a linear mixed-effect model. The model fits a nonlinear relationship by a restricted cubic spline with 4 knots at the 5th, 35th, 65th, and 95th percentiles of the data, when the condition (Clemastine versus vehicle) and condition-by-time interactions were performed as fixed effects. A likelihood ratio test to detect group-wise significant differences by non-overlapping 95% confidence intervals. Confidence intervals were calculated using case bootstrapping with 1000 iterations. 95% confidence intervals were computed as 1.96 times the standard error of the model fits. For immunohistochemical data, a two-way ANOVA (p < 0.05) was applied to determine the significances in fluorescent markers between Clemastine and vehicle tissue. The Fisher’s Least Significant Difference (LSD) was applied to identify group-wise significant differences in the implant side between Clemastine and vehicle conditions. The Dunnett’s test was used to compare the intensity/cell density at each bin in the implant side to the contralateral controls in each condition (Clemastine, vehicle). Unequal-variance Welch’s t-test was applied to detect any significance in contralateral histological metrics between two conditions.

## 3. Results

Clemastine is an antimuscarinic compound being investigated to treat demyelinating neurodegenerative diseases [56, 62, 82, 83, 94]. Administration of Clemastine effectively promotes OPC differentiation, enhances myelination, and rescues severe symptoms of inflammatory demyelination disorders [56, 94–96]. To determine the impact of Clemastine on electrophysiological recording performance over the chronic timescale, C57BL/6J wildtype mice were treated with Clemastine (Fig. 1A). In order to maximize the effect of Clemastine, subjects were pre-conditioned 7 days prior to the surgery, and then continued daily for 16 weeks following the intracortical microelectrode implantation. The dosage of Clemastine was determined based on literature [55, 56, 62, 79, 95]: Clemastine was dissolved into 10% DMSO/sterile PBS vehicle solution and administered at a dose of 10 mg/kg via intraperitoneal injection every day. Vehicle-only solution was injected into the control animal in the same delivery manner to control for the impact of the delivery method on recording performance. The rationale of pre-conditioning was that Clemastine has been shown to mitigate behavioral symptoms of neurological disorders with consecutive 7 days administration [62, 79, 83]. Specifically, these studies observed increased OPC differentiation to mature myelinating oligodendrocytes and remyelination in a hypoxia-induced neuroinflammatory environment similar to implantation-induced inflammatory cascades [62]. Therefore, we expected that the pre-treatment with Clemastine would enhance OL differentiation in the acute inflammation stage following microelectrode implantation and improve recording performance during acute to ‘early chronic’ period.

**Figure 1:**
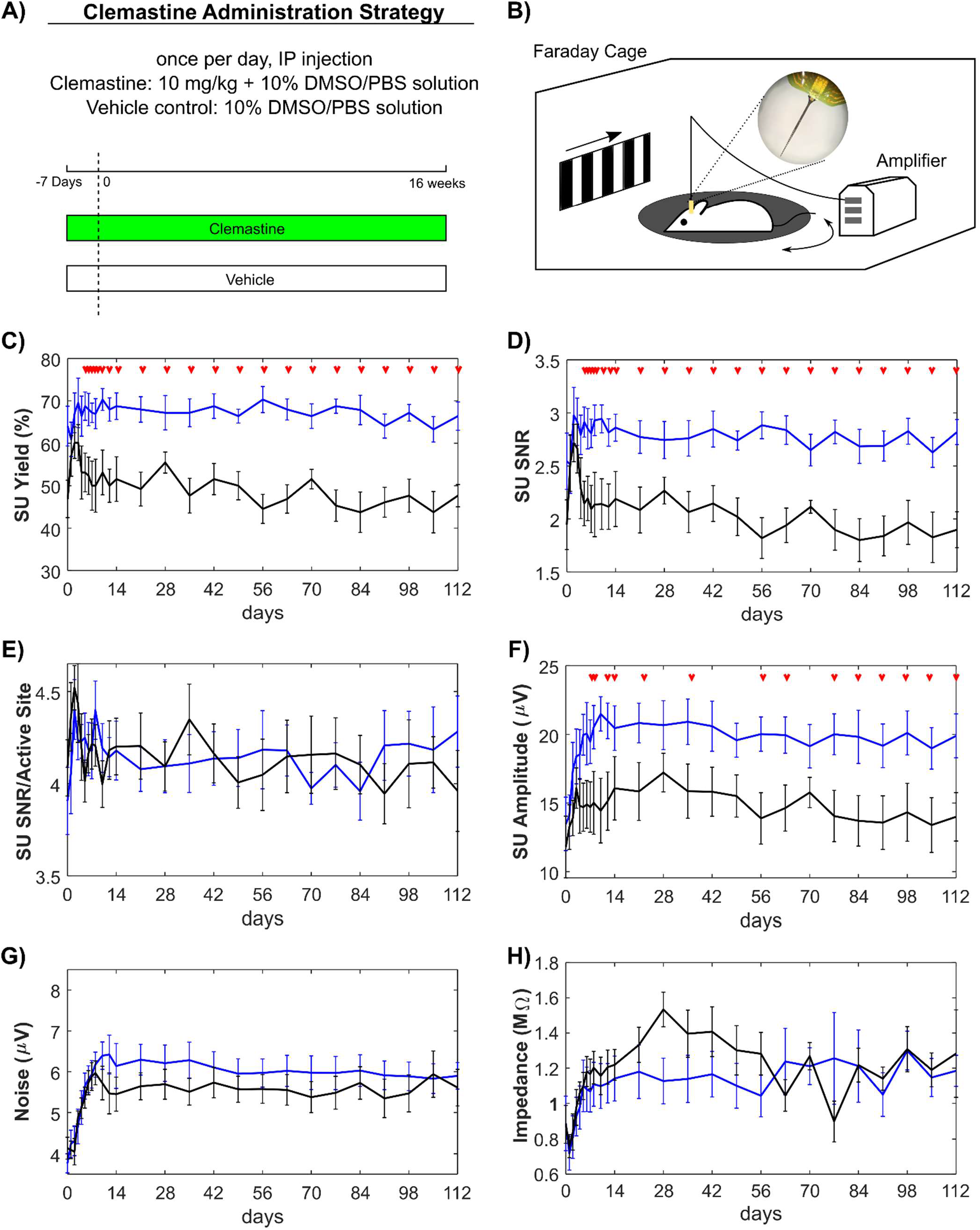
Clemastine improves single-unit (SU) activity of the implanted microelectrodes over chronic 16 weeks. A) Schematic illustration of Clemastine administration strategy and timetable in experimental and control groups. Mice received daily Clemastine (10 mg/kg dissolved in 10% DMSO/PBS solution) or vehicle solution (10% DMSO/PBS solution) via intraperitoneal (IP) injection from 7 days prior to microelectrode implantation until 16 weeks post-implantation. Day 0 indicates the day of microelectrode implantation surgery. B) Electrophysiological recording setup. Mice were awake, head-fixed on a rotating platform in a Faraday cage. The drifting bar gradient on LCD monitor was applied as the visual stimulation paradigm. Clemastine treated group (blue) and vehicle only control (black) were plotted over time for the following metrics; Clemastine administration led to significantly robust SU yield (C), signal-to-noise ratio (SNR) (D), and signal amplitude (F), while the SNR over active sites only (E), noise floor (G), site impedances at 1 kHz (H) were statistically comparable with vehicle group. Red arrows indicate significant differences between Clemastine-treated and vehicle control mice by a linear mixed-effects model following likelihood ratio test with a 95% confidence interval.

### 3.1 Clemastine effectively enhances functional recording performance

#### 3.1.1 Clemastine improves neuronal single-unit activity throughout the 16-week implantation in a depth-dependent manner

To examine Clemastine’s influence on the functionality of the implanted microelectrode, electrophysiological data was recorded from awake, head-fixed mice inside an electrically grounded Faraday cage. The cage was enclosed in a dark room for spontaneous, resting recording sessions, while a drifting bar gradient was presented on a monitor to the contralateral eye for visual evoked recording sessions (Fig. 1B). We first compared the single-unit (SU) recording metrics between Clemastine and vehicle groups independent of laminar depth and averaged all channels along the laminar shank of the microelectrode. Clemastine administration was hypothesized to increase the SU recording quality for chronically implanted microelectrode compared to vehicle controls, with increased SU availability and signal amplitude. SU yield was calculated as the percent of channels on the array that detected at least one SU, which can be used to measure the availability of neuronal SU sources near the chronically implanted microelectrode. Note that not all channels are expected to detect single-units because one channel usually ends up in layer 1 and 3-4 channels end up in the collosum cassette, where there are no neuronal somas.

The SU yield (66.41% ± 9.41%) in Clemastine-treated mice was significantly higher than the SU yield in vehicle controls (47.66% ± 7.42%) at the end of 16-week implantation (Fig. 1C, *p* < 0.0001). The significant difference in SU yield between Clemastine and vehicle groups occurred at day 6 post-implantation (non-overlapping 95% confidence intervals). While SU yield of Clemastine-treated group was relatively stable throughout the 16-week implantation (day 0: 64.06% ± 12.50%), vehicle mice declined over time by approximately 10 %. Similarly, the SU signal-to-noise ratio (SNR), which measures the strength of the SU activity, was significantly higher (Fig. 1D, p < 0.0001) in Clemastine mice compared to vehicle group starting from day 5 post-implantation (likelihood ratio test, non-overlapping 95% confidence intervals). The averaged SU SNR in Clemastine mice was 2.54 ± 0.75 at day 0 post-implantation, which was similar to 2.82 ± 0.34 by 16-week administration. In contrast, the averaged SNR in vehicle mice was reduced from 2.53 ± 0.76 on day 0 to 1.90 ± 0.48 on week 16. Furthermore, the strength of individual SU quantified as the averaged SNR in only active recording sites showed no significant difference between Clemastine and vehicle mice (Fig. 1E, *p* = 0.9631). Together, these results suggest that the Clemastine increases availability of neuronal sources rather than changing firing properties of surviving neurons.

Besides SNR, the strength of SU action potential showed significant elevation in the average amplitude of the SU waveform of the Clemastine group (19.89 μV ± 4.52 μV) compared to that of the vehicle group (13.99 μV ± 4.98 μV) starting at day 6 post-implantation (Fig. 1F, p < 0.0001). The noise profile of the two groups were comparable (Fig. 1G, Clemastine: 6.34 μV ± 0.93 μV, vehicle: 6.47 μV ± 0.98 μV), with both groups experiencing low noise fluctuations before stabilizing to a steady level after approximately 2 weeks. Similarly, the device impedance experienced fluctuations and then stabilized starting day 14 post-implantation (Fig. 1H, Clemastine: 1.14 ± 0.36 MOhms, vehicle: 1.22 ± 0.25 MOhms). There was no significant difference in device impedance between the Clemastine and vehicle groups throughout the 16-week implantation (p = 0.5689). The statistical comparable patterns in noise and impedance between Clemastine and vehicle groups suggest that the Clemastine has less influence on electrophysiological properties of environmental background or glial scar insulation in tissue near the implanted microelectrodes.

To determine how Clemastine improves the detectability of SU activity longitudinally at different depths, we examined the cortical layer dependent differences in SU activity between the Clemastine-treated group and vehicle control. The depth was aligned to layer IV for each animal and each time point by using visually evoked current source density as previously described [84]. Briefly, Layer IV depth location was identified as the first inward current in visual cortex. Then, the SU recording metrics were plotted along the aligned depth over the implantation time as heatmaps to examine the regionspecific effect of Clemastine on cortical and hippocampal CA1 microelectrode recording performance. For SU yield (Fig. 2A), Clemastine-treated mice maintained significantly elevated SU yield in cortex relative to vehicle control starting day 12 post-implantation (Fig. 2B, Clemastine: 71.88% ± 27.82%, vehicle: 57.81% ± 16.28%; p < 0.05). In hippocampus, Clemastine group had significant higher SU yield in CA1 from 6 to 9 weeks post-implantation before the SU Yield declined to the level of vehicle controls (Fig. 2C, p = 0.08845, likelihood ratio test with non-overlapping 95% confidence intervals), suggesting that the reduction in SU yield in CA1 was delayed in Clemastine mice.

**Figure 2:**
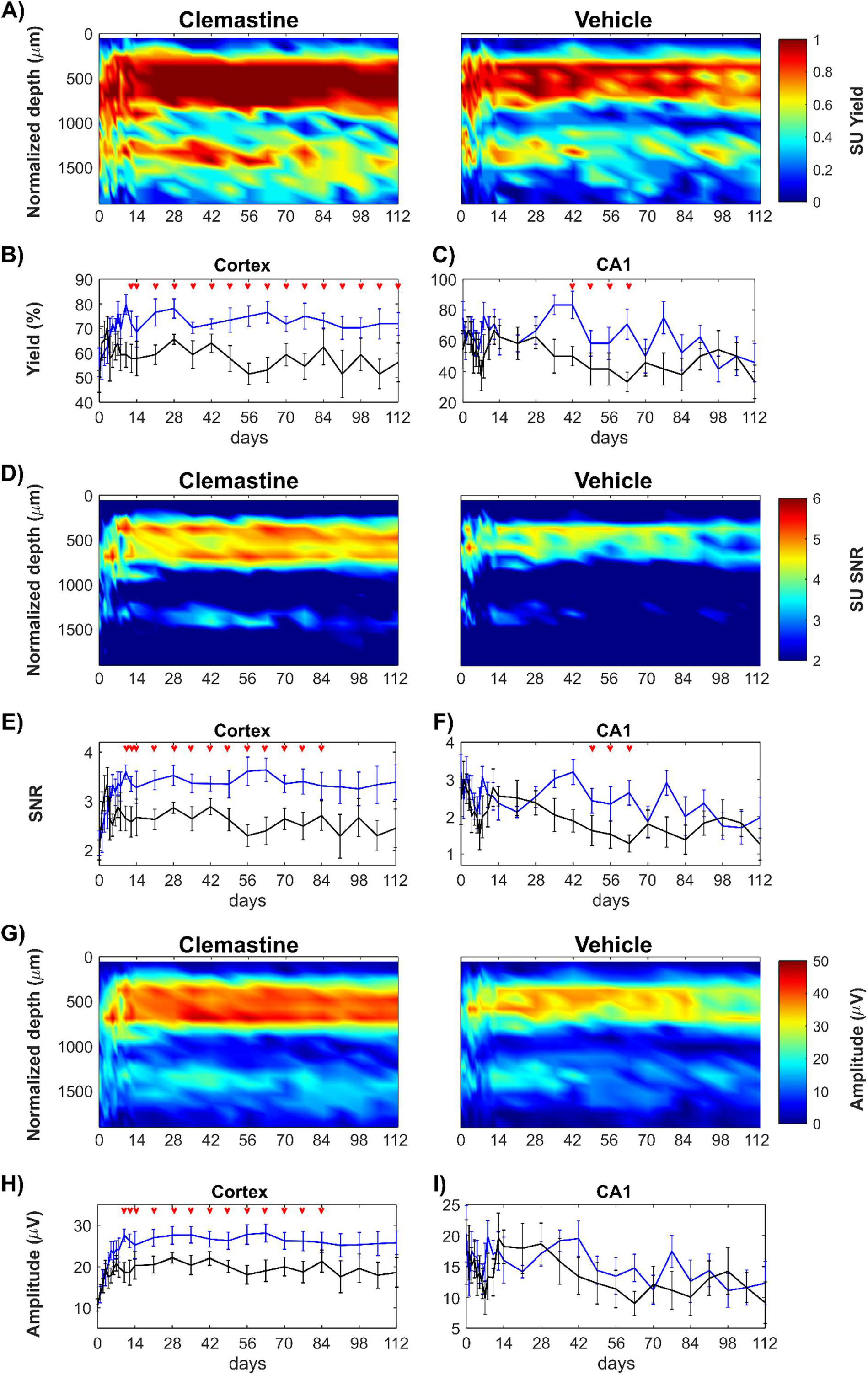
Clemastine increases SU recording quality of chronically implanted microelectrode effects on both cortical and hippocampal regions. The electrophysiological metrics of SU activity is plotted as a function of time and depth for Clemastine-treated mice (blue) and vehicle controls (black): Clemastine administration increases SU yield in a depth dependent manner relative to vehicle controls over chronic microelectrode implantation (A). SU yield averaged in cortical region (B) and hippocampus CA1 region (C) shows Clemastine’s positive effect is depth dependent. SU SNR as a function of depth and days post-implantation is plotted for Clemastine and vehicle mice (D). The SU SNR was averaged in cortical regions (E) and hippocampus CA1 region (F), respectively. SU amplitude over depth and implantation time was plotted as heatmaps for both Clemastine and vehicle animals (G). The cortical SU amplitude remained robust in Clemastine animal over time (H), while the signal amplitude in hippocampal CA1 region (I) was comparable between Clemastine and vehicle groups. Red arrows indicate non-overlapping 95% confidence intervals between Clemastine-treated and vehicle control mice at each time point by likelihood ratio test.

Similary, SU SNR (Fig. 2D) was plotted in along cortical depth over time. The cortical SNR was significantly higher in Clemastine-treated mice compared to vehicle controls (Fig. 2E, Clemastine: 3.28 ± 0.93, vehicle: 2.67 ± 1.06; p < 0.05, likelihood ratio test). In contrast, the hippocampal CA1 SNR experienced gradual decline in both Clemastine and vehicle groups. However, Clemastine group maintained the significantly higher SNR at 2.64 ± 0.93 at 9 weeks post-implantation compared to vehicle control that had a lower SNR level of 1.28 ± 0.64 (Fig. 2F, p = 0.1648, likelihood ratio test with nonoverlapping 95% confidence intervals).

Cortical SU signal amplitude was significantly higher in Clemastine group compared to vehicle control, specifically 2-11 weeks post-implantation (Fig. 2H, p < 0.05). However, there was no significant difference in SU signal amplitude in hippocampal CA1 between Clemastine and vehicle groups (Fig. 2I, p = 0.7922). Both groups experienced reduction in signal amplitude over the chronic implantation period, declining nearly by 39% in Clemastine group and 47% in vehicle controls, respectively. The depth profile of noise floor demonstrated that both Clemastine and vehicle groups had comparable performance in cortical and hippocampus CA1 regions by 16-weeks post-implantation (Supplementary Fig. 1, cortex: p = 0.2444; hippocampus CA1: p = 0.5166).

To summarize, Clematine-induced neuroprotection that is region-specific. Administration of this promyelinating drug largely increased the cortical SU recording performance throughout the 16-week chronic microelectrode implantation period. In contrast in hippocampus CA1, Clemastine only delayed the loss of SU signal until 10 weeks post-implantation.

**Supplementary Figure 1:**
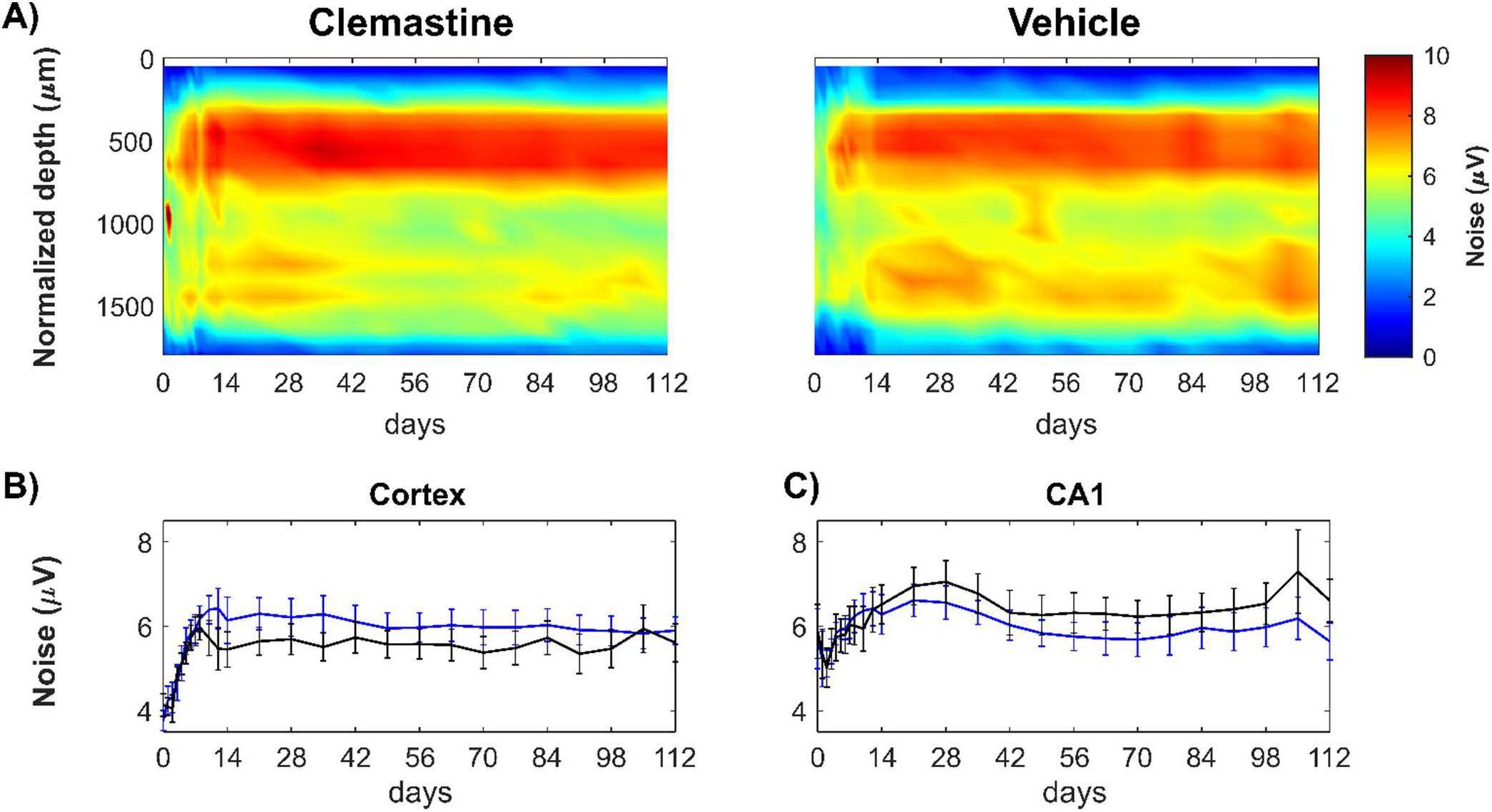
Noise floor between Clemastine and vehicle groups in cortex and hippocampus CA1, respectively. A) Noise floor between Clemastine-treated and vehicle mice is plotted as a functional of time and normalized depth. The noise floor was comparable between Clemastine and vehicle mice in cortex (B) and hippocampal CA1 (C).

#### 3.1.2 Clemastine rescues the deficits in populational neuronal firing activity

As Clemastine administration effectively improved SU activity throughout the 16 week implantation period in a depth-dependent manner, we next investigated the effects of Clemastine on the functional neural activity, which we measured as MU firing rate changes during spontaneous and visual evoked recording sessions. To examine whether Clemastine improved neuronal population activity during resting state as well as a functinonal network activity in response to visual stimuli, we sorted MU as all threshold-crossing events and plotted the firing rate over time in a depth-dependent manner (Fig. 3A, 3B). In the cortex, the average MU firing rate in Clemastine-treated mice was significantly higher than in vehicle controls from 13-16 weeks post-implantation for both resting state and visual evoked activty (Fig. 3C, 3E, linear mixed model following likelihood ratio test with non-overlapping 95% confidence intervals).

**Figure 3:**
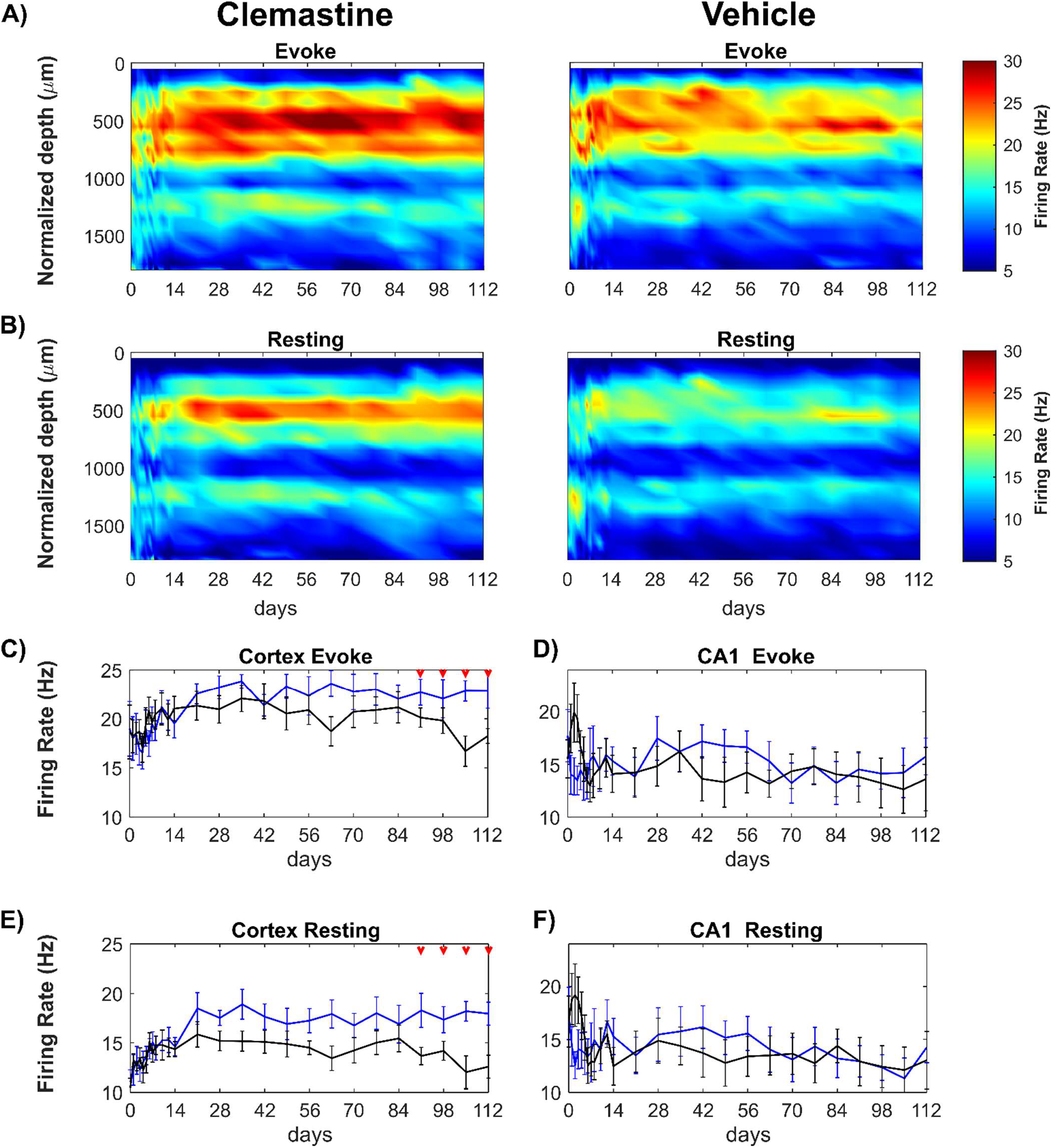
Clemastine leads to an improved functional activity compared to vehicle controls in a time- and region-dependent manner. Average multi-unit firing rate was plotted over time and aligned depth as heatmap during visual stimulation (A) and resting state (B) between Clemastine and vehicle mice. The multi-firing rates was plotted in region- and condition-specific manner for Clemastine (blue) and Vehicle (black) mice. Clemastine administration prevented the loss of the cortical MU firing rate at chronic 13-16 weeks post-implantation during visual stimulation (C) as well as resting state (E). However, there was little difference in hippocampal CA1 MU firing rate between Clemastine and Vehicle mice during either visual stimulation (D) or resting state (F), although Clemastine-treated mice had a low trend in hippocampal CA1 MU firing rate during acute 7 days post-implantation relative to vehicle controls. Red arrows indicate as significant difference between Clemastine-treated and vehicle control mice by a linear mixed-effects model following by likelihood ratio test with a 95% confidence interval.

Although both groups experienced fluctuation in MU firing rate during the initial 2 weeks post-implantation, the Clemastine group maintained a steady MU firing rate during visual stimulation over the 16 weeks implantation period (Fig. 3C). However, the visual evoked MU firing rate of the vehicle control group was reduced by nearly 13% to 18.24 ± 2.14 Hz by 16 weeks post-implantation (Fig. 3C). Similarly, the MU firing rate during resting-state remained elevated in the Clemastine group (17.96 ± 3.08 Hz) but the vehicle control group declined by approximately 12 % over the same 16 wk period (12.58 ± 3.30 Hz at 16 weeks post-implantation). This resulted in statistically significant decrease from 13 to 16 weeks post-implantation (Fig. 3E, p < 0.05, likelihood ratio test). Additionally, the visual stimulation consistently resulted in elevated cortical MU firing rates in both Clemastine and vehicle animals over the chronic 16-week implantation (Supplementary Fig. 2A, 2B), which indicates that recorded multi-units are functionaly integrated in the neural network in cortex. Together, these analyses suggest that Clemastine treatment preserves the quality of neuronal population firing activity, preventing decline in neuronal functionality in cortex at chronic timepoints.

Having demonstrated that Clemastine contributes to preservation of functional neural activity in the cortex over the chronic period (13-16 wks), we investigated the impact of Clemastine on functional neural recording in CA1 hippocampus (Fig. 3D, 3F). Interestingly, there were no significant differences in MU firing rate in hippcampus CA1 between Clemastine and vehicle groups over 16-week implantation, neither during visual stimulation (Fig. 3D, p = 0.3700), nor during resting state (Fig. 3F, p = 0.2135). Although cortical MU firing rate increased approximately 27% during stimulation compared to resting-state, the changes in MU firng rate in hippocampus CA1 was only about 3% for both groups (Supplementary Fig. 2C, 2D). This small increase in viaully evoked MU firing rate in hippocampus CA1 suggests there is limited activation of CA1 neurons during drifting bar gradient paradigm. Interestingly, the vehicle mice exhibited a peak in hippocampus CA1 MU firing (19.93 ± 8.02 Hz) at 3 days post-implantation. However, this peak was not observed in Clemastine group, which had average MU firing rate at 15.33 ± 3.53 Hz during initial 7 days post-implantation. This distinct firing rate in hippocampus CA1 between Clemastine and vehicle mice may indicate that Clemastine could provide neuroprotection to avoid hyperexcitability in hippocampus CA1 region (Fig. 3D). Together, these analyses suggest that Clemastine treatment preserves the quality of populational neuronal firing activity, preventing decline in neuronal functionality in cortex at chronic timescale, and protecting potential hyperexcibility in hippocampus CA1 during the acute neuroinflammation period.

**Supplementary Figure 2:**
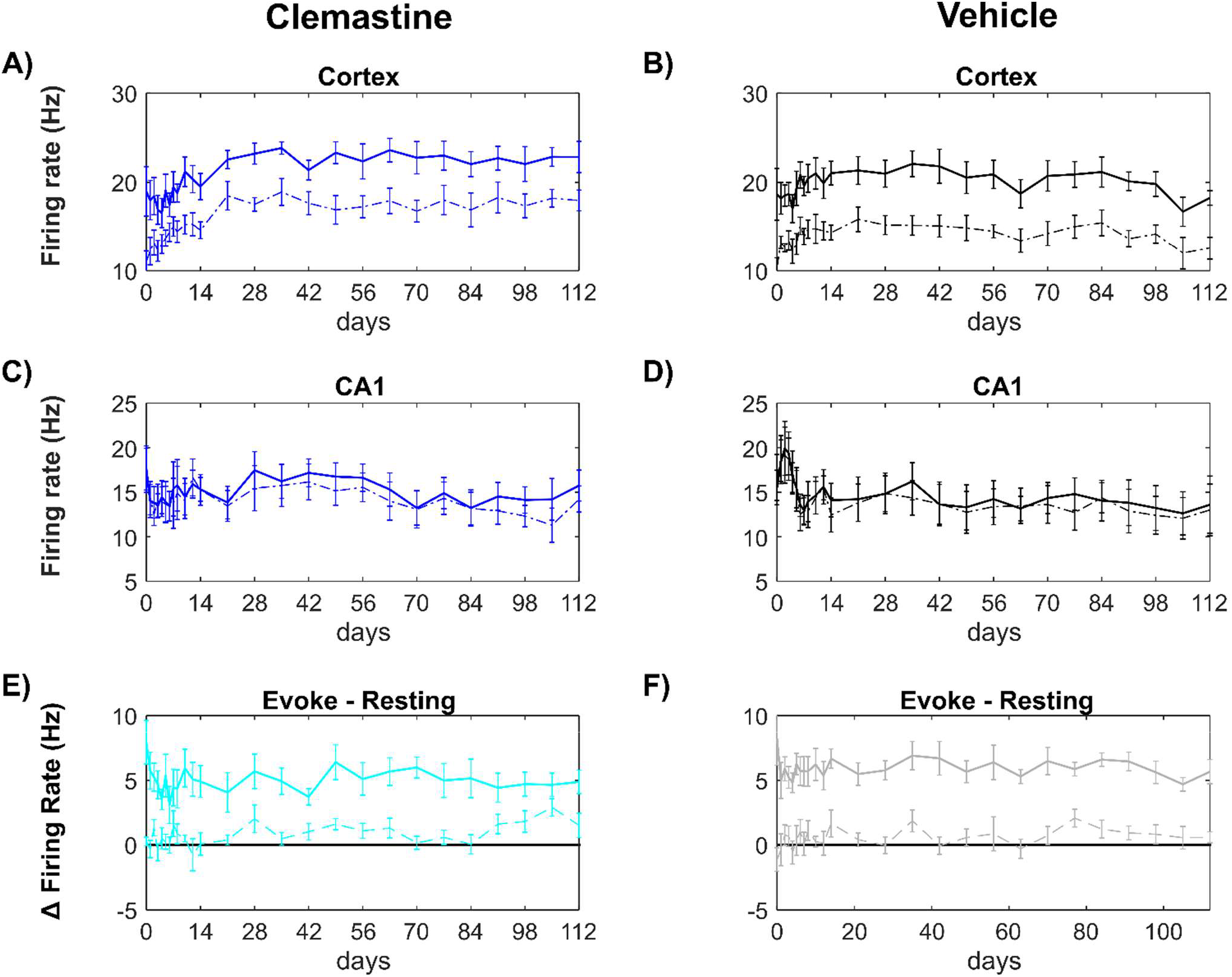
The increases in MU firing rate by visual stimulation was significant in cortex but was comparable to resting state in hippocampus CA1 in both Clemastine and vehicle mice. MU firing rate during visual stimulation (solid lines) was elevated relative to resting state (dash lines) by nearly 27% in cortex (averaged across all animals). Clemastine (A) and vehicle (B) mice increased firing rate of visual evoke compared to resting state over the entire 16 weeks implantation. The MU firing rate in hippocampus CA1 was about 3% increase in visual stimulation compared to resting state conditions. Both Clemastine (C) and vehicle (D) animals exhibited comparable firing rate between visual evoke and resting conditions over 16 weeks implantation. The change in firing rate between evoke and resting state in cortex (solid lines) and hippocampus (dash lines) were plotted in Clemastine (E) and vehicle (F) conditions.

#### 3.1.3 Clemastine prevents the chronic loss of functional oscillatory activity in a frequency-specific manner

The SU and MU analyses showed that enhancing OL and myelin activity with Clemastine improves neural activity in the microenvironment near the implant (less than 80~160 μm [97]) over the chronic implantation period. Therefore, we next asked how Clemastine alters neural oscillatory activity, which is related to neuronal population activity over long distances. Local field potential (LFP) measured during visual stimulation was used to evaluate functional oscillatory activity in visual cortex. The impact of Clemastine administration was examined by power spectral analysis of LFP activity between 0.4-300 Hz relative to vehicle controls.

To determine if Clemastine administration affects LFP oscillations during functionally evoked network activity, normalized evoked power was quantified as the changes in power during visual stimulation relative to resting state and was evaluated over time and with respect to laminar depth and specific frequency bands. We found that both Clemastine and vehicle treated mice demonstrated that most of the evoked oscillatory activity was on lower frequency bands, approximately 0.4-30 Hz (Fig. 4A, 4B). The depth profile showed that normalized evoked power in both groups was primarily located at ~50 - ~900 μm below the surface, which corresponds to cortical depths (Fig. 4C). However, the heatmaps showed a reduction in normalized evoked power of vehicle controls over time (Fig. 4C). Therefore, we first averaged the normalized evoked power over the entire frequency range (125 Hz) to examine whether there were significant differences in mean power between Clemastine mice and vehicle controls (Fig. 4D, *p* = 0.1451). Over 14-16 weeks post-implantation, the vehicle control group experienced a significant decline in mean power (nearly 42%) whereas the Clemastine group remained stable (Fig. 4D, likelihood ratio test with non-overlapping 95%confidence intervals).

**Figure 4:**
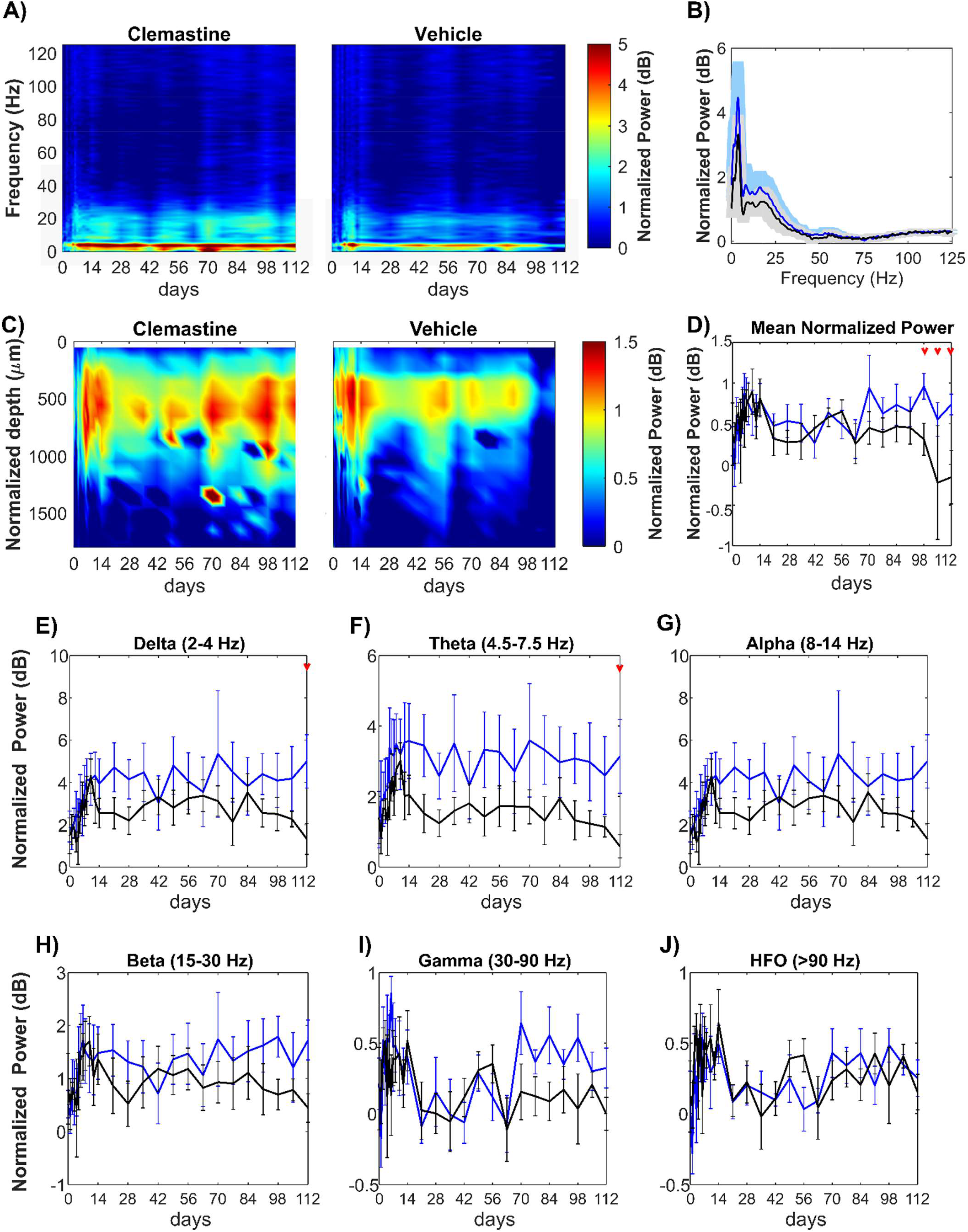
Clemastine effects on oscillatory activity during visual stimulation at specific frequency range. (A) Heatmaps of average evoked power normalized to spontaneous power as a function of frequency and implantation time for Clemastine and vehicle groups. (B) Spectral distribution of normalized evoked power over 0.4 - 125 Hz frequencies for Clemastine (blue) and vehicle (black) animals. (C) Heatmaps of the average normalized visually evoked power plotted as a function of aligned cortical depth and implantation time. Cortical regions had an increased normalized evoked power relative to hippocampus CA1. (D) Mean LFP power over 0.4 −300 Hz between Clemastine and vehicle groups over time. Clemastine administration prevented the loss of mean power compared to the vehicle group from 14-16 weeks post-implantation. Normalized visually evoked power at individual frequency bands, delta (E), theta (F), alpha (G), beta(H), gamma (I), high frequency oscillation (HFO) (J), between Clemastine and vehicle groups over time. Red arrows indicate significant differences between Clemastine-treated and vehicle control mice by a linear mixed-effects model following by likelihood ratio test with a 95% confidence interval.

Next, different frequency bands of the normalized evoked power of Clemastine- and vehicle-treated mice were examined over time. Across delta (2-4 Hz) and theta (4.5-7.5 Hz) frequency ranges, Clemastine-treated mice exhibited more stable power throughout the 16-week implantation period. In contrast, the vehicle control group experienced a gradual decline over 14-16 weeks post-implantation, resulting significant differences between two groups at 16 weeks post-implantation (Fig. 4E, delta band: p = 0.1694; Fig. 4F, theta band: p = 0.0725, likelihood ratio test with non-overlapping 95% confidence intervals). Similarly, frequency oscillation below 2 Hz showed separation of the power spectrum between Clemastine and vehicle groups (Supplementary Fig. 3A). Specifically, the power of vehicle control mice significantly dropped relative to Clemastine mice at week 16 post-implantation (Supplementary Fig. 3B, likelihood ratio test with nonoverlapping 95% confidence intervals). There were no significant differences in normalized evoked power between Clemastine and vehicle mice over alpha (8-14 Hz, Fig. 4G), beta (15-30 Hz, Fig. 4H), gamma (30-90 Hz, Fig. 4I) and high frequency oscillation (HFO, > 90 Hz, Fig, 4J). However, Clemastine mice showed a substantial increase in gamma power (30-90 Hz) compared to vehicle mice from around 10 weeks post-implantation (Fig. 4I, p = 0.2947). Overall, the results of LFP analysis indicate that Clemastine helps sustain the oscillatory activity in visually evoked visual cortex over long-term microelectrode implantation, especially in low frequency delta, theta bands.

**Supplementary Figure 3:**
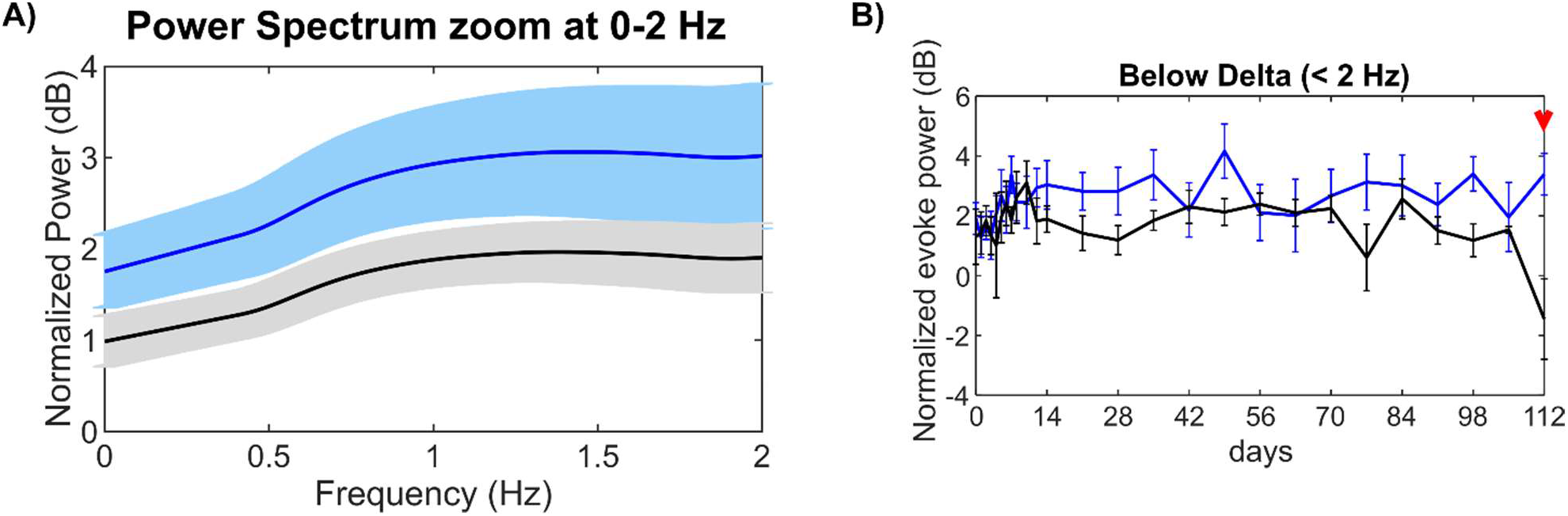
Clemastine affects slow oscillatory activity below 2Hz during visual stimulation. (A) Normalized evoke power spectra below 2 Hz for Clemastine (blue) and vehicle (black) groups. (B) Normalized evoke LFP power below 2 Hz. Clemastine (blue) significantly rescued the drop that occurred in vehicle condition (black) at 16 weeks. Red arrow indicates significant difference between Clemastine-treated and vehicle control mice by a linear mixed-effects model following by likelihood ratio test with a 95% confidence interval.

### 3.2 Clemastine alters network circuit near the implanted microelectrode

#### 3.2.1 Clemastine improves feasibility and functionality of putative neuronal subtypes over the chronic implantation period

After demonstrating that Clemastine administration improves SU, MU, and LFP functional recording performance with chronically implanted microelectrodes, we next asked whether this improved detection of neuronal activity resulted from Clemastine’s effect on neuronal subtypes and functional circuits near the microelectrode. To investigate how different neuronal subtypes are recruited for processing of visual evoked information between Clemastine and vehicle mice, the recorded single-units were classified by the action potential waveform width, which is the trough-to-peak latency of the waveform [88, 89] (Supplementary Fig. 3). The bimodal distribution of SU waveform widths revealed two distinct populations of the waveforms: the putative inhibitory neurons with narrow waveform peaked at 0.25 ms, and the putative excitatory neurons with wide waveform peaked at 0.54 ms (Supplementary Fig. 3). Thresholding at 0.41 ms yielded 8.55% putative inhibitory neurons and 91.45% putative excitatory neurons of the 8736 total isolated SU action potentials.

Then, the detectability of putative inhibitory and excitatory neurons was examined in the form of average yield of putatively classified subtypes over time and depth (Fig. 5A, 5B). More putative excitatory neurons were detected in Clemastine-treated mice in both cortical and hippocampal regions compared to vehicle controls (Fig. 5A). Additionally, the Clemastine group maintained a higher yield in putative inhibitory neurons in cortical L2/3, L4, and L5/6 relative to vehicle group (Fig. 5B). The average number of putative excitatory neurons for each microelectrode array was significantly higher in Clemastine treated mice compared to vehicle controls starting 5 weeks post-implantation and throughout the whole recording period (Fig. 5B, p < 0.05). Meanwhile, putative inhibitory neuron recording viability was significantly higher in Clemastine-treated mice compared to vehicle controls, especially between 11-13 weeks post-implantation (Fig. 5D, *p* =0.1559, likelihood ratio test, non-overlapping 95% confidence intervals). These results indicate that Clemastine increased the detectability of putative excitatory and inhibitory neuronal subtype near the implanted microelectrode.

**Figure 5:**
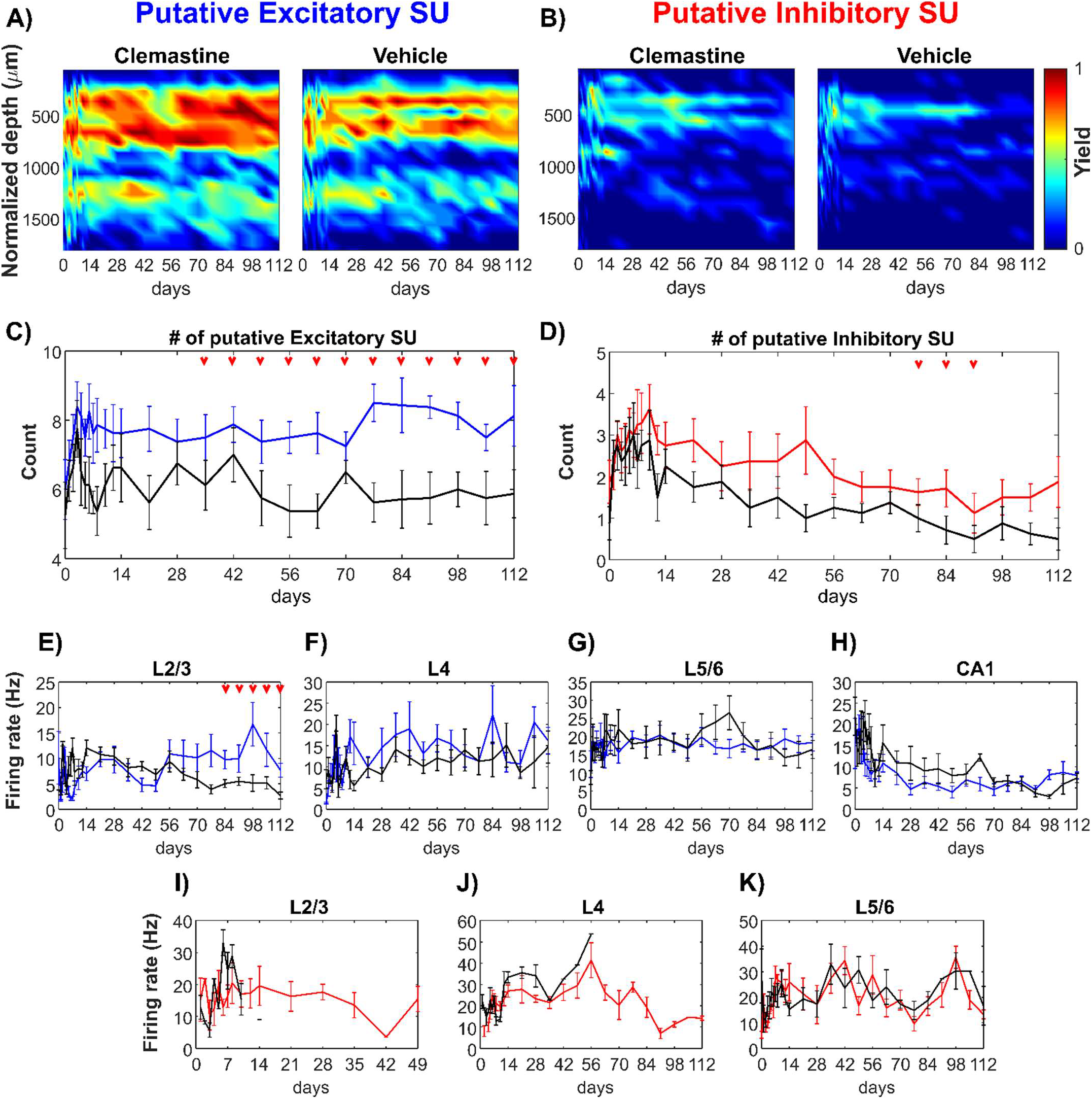
Clemastine enhances putative neuronal subtype detectability and functionality over chronic implantation. Yield heatmaps for putative excitatory (A) and inhibitory neurons (C) plotted as a function of time and depth between Clemastine and vehicle groups. The number of putative excitatory (B) and inhibitory neurons (D) across all 16 channels between Clemastine-treated mice (colored-lines) and vehicle controls (black lines) over chronic 16-week implantation period. Clemastine administration increased the yield of both putative excitatory and inhibitory neurons relative to vehicle group. The visual evoke firing rates of putative excitatory neurons in L2/3 (E), L4 (F), L5/6 (G), and hippocampus CA1 (H) between Clemastine- and vehicle-treated mice over chronic implantation. Further, Clemastine’s effect on putative inhibitory neuron firing rate was compared to the control group in L2/3 (I), L4 (J), and L5/6 (K) relative to vehicle controls. IF there was no putative inhibitory neuron detection, the firing rate would be NA instead of zero. Therefore, the loss of putative inhibitory neuron detection in L2/3 and L4 resulted in loss of putative inhibitory firing rate measurements. Red arrows indicate significant differences between Clemastine-treated and vehicle control mice by a linear mixed-effects model following by likelihood ratio test with a 95% confidence interval.

Next, the impact of Clemastine on putative neuronal subtype functionality was examined through differences in evoked firing rate between Clemastine and vehicle groups at different cortical depths. After the initial two-week fluctuation, L2/3 putative excitatory neurons in vehicle mice showed a gradual reduction in firing rate over time (Fig. 5E, 12.08 ± 4.35 Hz at 2 weeks post-implantation to 2.72 ± 1.97 Hz at 16 weeks post-implantation). In contrast, in Clemastine-treated mice, the putative L2/3 excitatory firing rate were significantly greater compared to the vehicle controls between 12-16 weeks post-implantation (Fig. 5E, p < 0.05). In L4 (Fig. 5F), L5/6 (Fig. 5G), and hippocampal CA1 (Fig. 5H), the putative excitatory firing rates in Clemastine group were comparable to vehicle controls. The putative inhibitory firing rate demonstrated a comparable pattern between Clemastine and vehicle groups in L5/6 (Fig. 5K, p = 0.7388). However, vehicle control mice had fewer detectable putative inhibitory sources in L2/3 and no putative inhibitory activity was detected after 2 weeks post-implantation (Fig. 5I). In contrast, Clemastine treated mice had higher putative inhibitory single-units with a more stable L2/3 firing rate, approximately at 15.44 ± 11.27 Hz, over longer implantation periods up to week 7 post-implantation (Fig. 5I). Furthermore, in vehicle control groups, putative inhibitory neurons in L4 could no longer be detected 4 weeks post-implantation, whereas Clemastine groups maintained L4 inhibitory firing rate (19.32 ± 13.52 Hz) before declining at week 13 post-implantation (Fig. 5J). Putative inhibitory neurons were not detected in CA1 for either group. Taken together, these data suggest that Clemastine improves the viability and functional strength of distinct neuronal subtypes in visual cortex around the implanted microelectrode arrays.

**Supplementary Figure 3:**
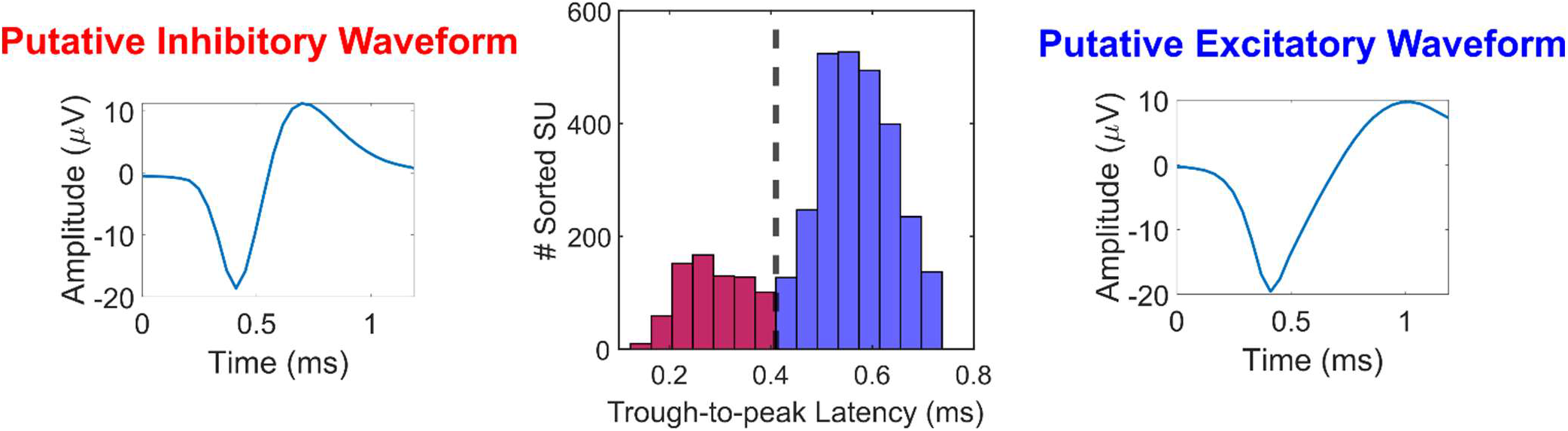
Neuronal functional subtype classification. Waveforms were classified based on action potential waveform trough-to-peak (TP) latency, following a bimodal distribution that separates putative inhibitory neurons (red, TP latency < 0.41 ms) and putative excitatory neurons (blue, TP latency ≥ 0.41 ms). Representative putative inhibitory waveform (left) and representative excitatory waveform (right).

#### 3.2.2 Clemastine modulates action potential transmission during local visual cortex activation

The previous analysis revealed that Clemastine improves the detectability and functional firing rates of putative neuronal subtypes during the chronic implantation period. Therefore, we then asked whether the responsiveness of neuronal population network activity to visual stimulus is influenced by the Clemastine administration. Responsiveness was characterized as MU yield and SNFRR, which were evaluated as the MUA within equal sized bins before and after visual stimulation, as previously described [90]. The calculation of MU yield and SNFRR depends on parameters such as bin sizes and latency from the stimulus onset. MU Yield was calculated as the percent of channels with significantly different MUA (p < 0.05) between the bins before and after the stimulus. SNFRR quantified the magnitude of the difference in MU firing rate between stimulus ON and OFF conditions (see Eq. 1 in Methods). The MU yield characterized the temporal patterns of MUA during 1-s visual stimulus ON period as a function of bin size and latency (Fig. 6A). Here, visual stimulation generated a strong, transient firing response, followed by a weaker sustained response, as previously observed in [75, 90]. However, because there are multiple combinations of bin sizes and latencies, it is important to fairly compare Clemastine and vehicle control for the same bin size and latency in order to investigate how Clemastine influences the temporal patterns of MUA responses during the 1-s visual stimulation period. To address this, we fixed the bin size to the value that optimize MU yield in the vehicle control condition averaged across all animals and all time points. The bin size that optimized MU yield in cortex was 46 ms and in hippocampus CA1 was 97 ms (Fig. 6A).

**Figure 6:**
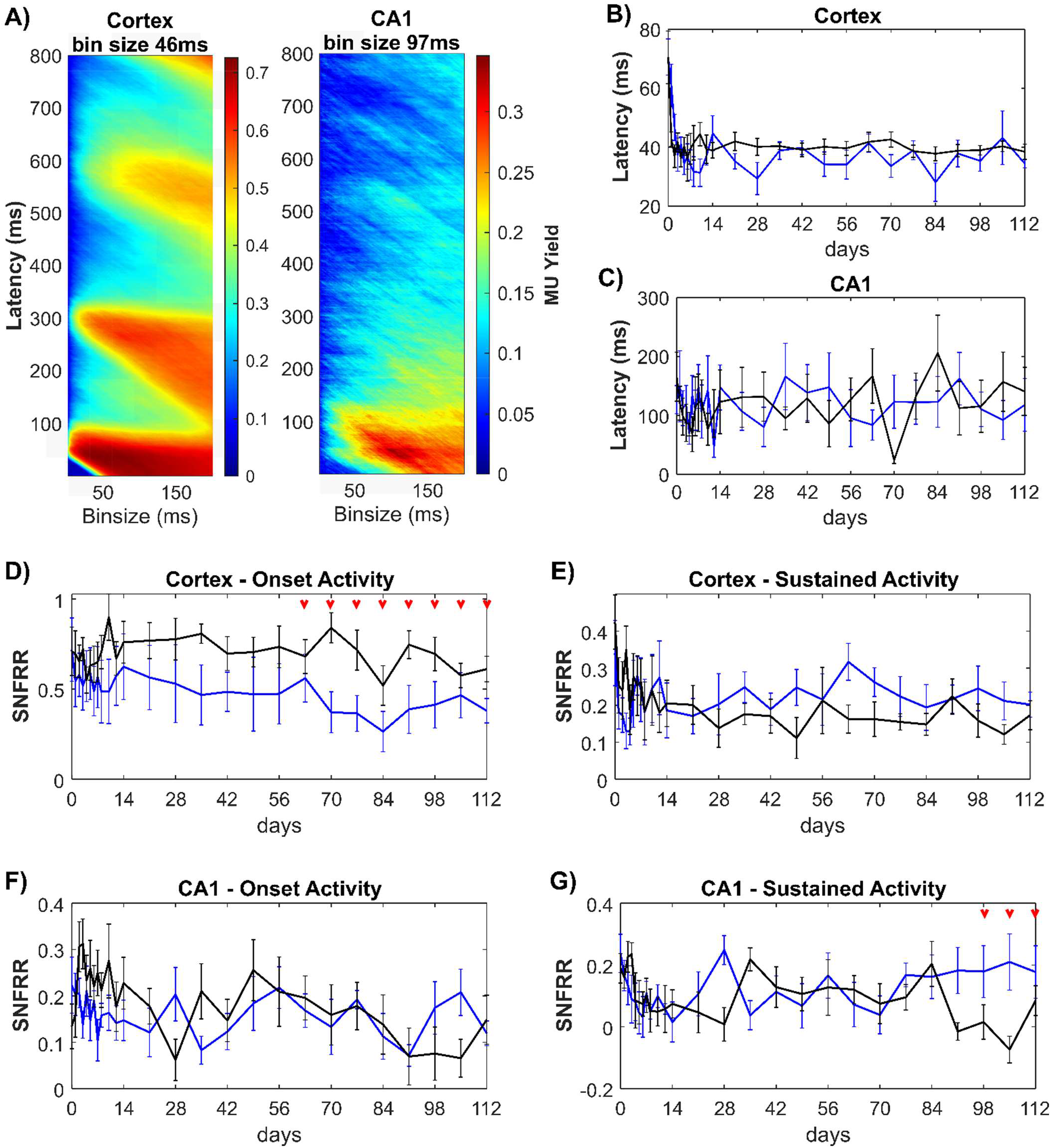
Clemastine modulates the functional activity of multiunit activity (MUA) over the chronic implantation period. (A) Averaged MU yield in the vehicle control group plotted across all time points dependent on bin sizes and latency in 1-ms resolution in response to 1-s ON stimulation period. There was a strong transient MUA within 0-100 ms, following by a weaker, sustained MUA from 100-800 ms. The bin sizes for optimal MU yield were 46 ms in cortex and 97 ms in hippocampal CA1 region, respectively. When the bin size was fixed, latency from the stimulus onset to the peak of MUA were comparable between Clemastine (blue) and vehicle (black) mice in cortex (B) and in hippocampus CA1 (C). The larger latency in hippocampus relative to cortex suggests visual information first reaches cortex and then the hippocampus. The transient MUA in response to visual stimulus onset was characterized as the averaged SNFRR within 100 ms in cortex (D) and hippocampus (F). The weaker, sustained or adapted MUA was quantified as the averaged SNFRR between 100-800 ms after stimulus onset in cortex (E) and hippocampus (F). SNFRR equals to zero indicates that there is no difference in MU firing rates before and after the visual stimulation. Red arrows indicate significant differences between Clemastine-treated and vehicle control mice by a linear mixed-effects model following by likelihood ratio test with a 95% confidence interval.

Since myelin can accelerate the action potential propagation along the axons [98], we asked whether the promyelinating Clemastine treatment resulted in shorter latency from the stimulus onset to the peak MU yield. However, there was no statistical differences in cortical latency between Clemastine and vehicle mice (Fig. 6B, p = 0.2780), which is in agreement with previous studies[75, 99]. The cortical latency in the Clemastine group was 34.5 ± 4.274 ms at 16 weeks postimplantation, which is similar to 38.5 ± 6.775 ms in the vehicle control group. Furthermore, there was no significant difference (p = 0.1809) in hippocampal latency between Clemastine (284.875 ± 34.235 ms) and vehicle (265.25 ± 45.049 ms) mice (Fig. 6C). These data indicate that Clemastine has no discernable effect on the latency of MUA response to visual stimulus.

As in Fig 6A, MUA exhibited a strong onset firing response and then a weaker sustained response during the 1-s ON visual stimulation period. We then asked whether Clemastine administration changes the MU firing rates during 0-100 ms onset and 100-800 ms sustained period. Averaged SNFRR during 0-100 ms showed that Clemastine mice had a significantly lower cortical SNFRR relative to vehicle control (Fig. 6D, p < 0.05), specifically 9-16 weeks post-implantation (likelihood ratio test, non-overlapping 95% confidence intervals). This difference suggests that Clemastine administration resulted in less MUA in cortex during the visual stimulation onset period in the microenvironment surrounding the electrode. Instead, there was no significance was detected in sustained cortical (100-800 ms) SNFRR (Fig. 6E, p = 0.4823). While depleting myelin resulted in significant changes in SNFRR at ~ 400 ms [58], we did not detect any significant difference in SNFRR during 350-450 ms between Clemastine and vehicle groups (data not shown). However, Clemastine-treated mice had an elevated trend in sustained cortical SNFRR compared to vehicle mice, suggesting Clemastine may increase MU firing rate during this period. In hippocampus, there was no difference in the onset SNFRR between Clemastine and vehicle groups over the 16 weeks implantation period (Fig. 6F, Clemastine: 0.1582 ± 0.0183; vehicle: 0.1913 ± 0.0212). However, vehicle mice experienced a loss of hippocampal sustained SNFRR (close to 0: 0.0849 ± 0.0170) over 13-15 weeks post-implantation, whereas the sustained SNFRR in Clemastine group was stable over time (0.1775 ± 0.0302). This difference in SNFRR was significant between two groups in this later timepoint (Fig. 6G, linear mixed model followed by likelihood ratio test, non-overlapping 95% confidence intervals). In general, Clemastine administration resulted a distinct profile in the functional responsiveness of population network activity near the implanted microelectrode, resulting in reduced onset MU firing in cortex and prevent the loss of sustained MU firing rate in hippocampus CA1.

#### 3.2.3 Clemastine enhances laminar connectivity along the implanted microelectrode

Since Clemastine influenced the action potential transmission in neural circuit near the microelectrode, we next examined whether Clemastine can also modulate functional connectivity between different laminar networks. Because Fig. 3D demonstrated limited recording of visual activity in hippocampus CA1, we focused on cortical functional connectivity between different layers by analyzing coherence. Coherence was used to measure the similarity of LFP activity between different pairs of electrode channels. Here, to determine the level of functional connectivity between different cortical depth, we measured the changes in coherence during visual stimulation relative to spontaneous session (Δcoherence) over frequency bands and laminar depth [100]. For L4 – L2/3 connectivity, we found an elevated Δcoherence over delta-theta bands (2-8 Hz) in Clemastine group rather than vehicle controls (Fig. 7A). Clemastine group maintained a positive Δcoherence over time, which means Clemastine mice experienced elevated functional connectivity between L4 and L2/3 by visual stimulus. However, the vehicle controls showed a negative Δcoherence (Fig. 7B), which indicates the L4 and L2/3 did not increase similarity in LFP oscillations during visual evoke activation. Additionally, over the course of last 13-16 weeks post-implantation, there was a slight increasing trend in L4-L2/3 Δcoherence in Clemastine mice, leading to significant difference compared to vehicle mice (Fig. 7B, p = 0.1076, likelihood ratio test with non-overlapping 95% confidence intervals). These data suggest that Clemastine increases the functional connectivity between L4 as visual input layer and L2/3 as information processing layer relative to vehicle control.

**Figure 7:**
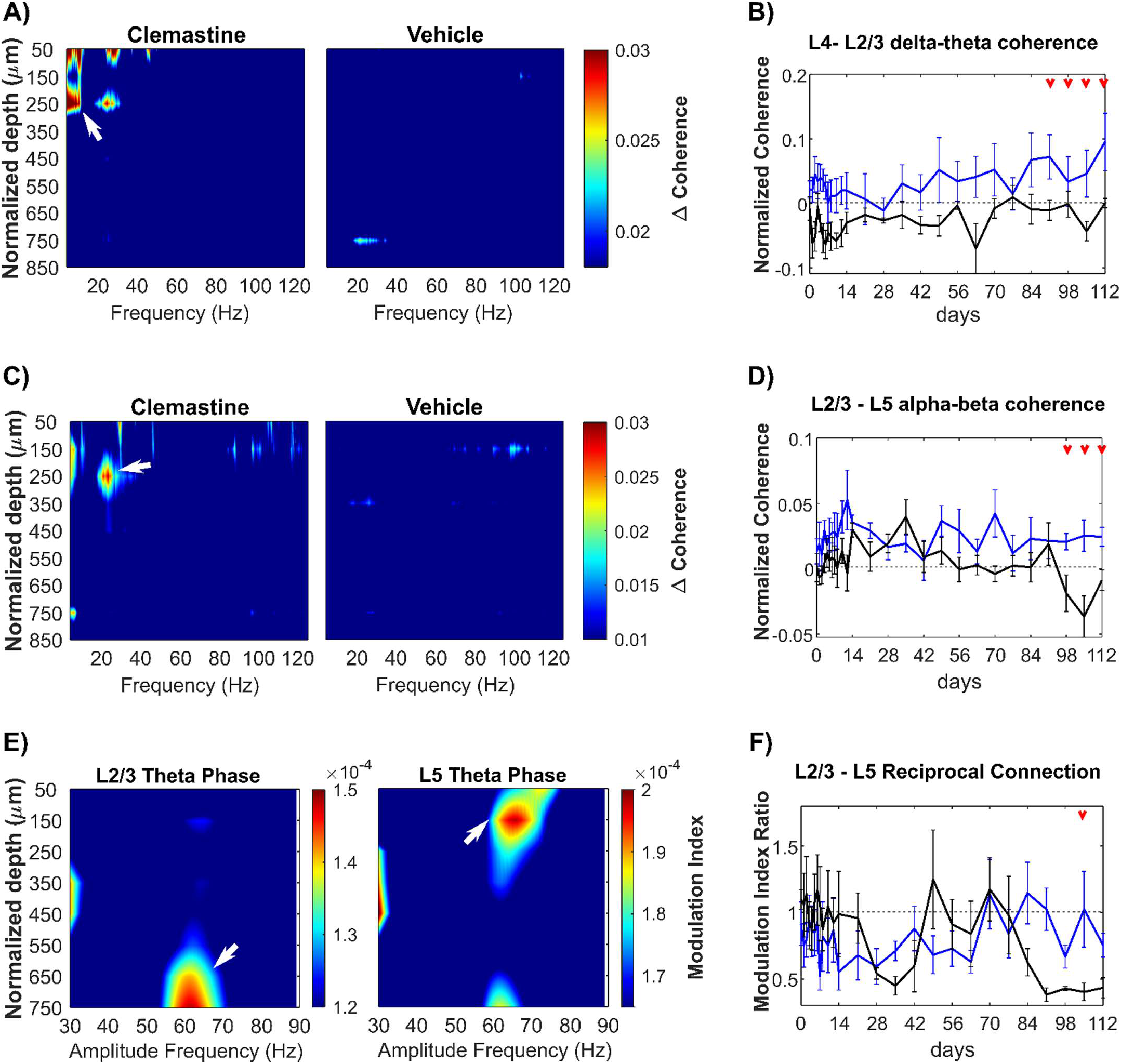
Clemastine improves the interlaminar functional connectivity over the chronic microelectrode implantation period. (A) Heatmap of normalized visually evoked coherence (Δcoherence) over depth paired with L4 electrode site as a function of frequencies. There was a robust Δcoherence (white arrow) between L4 and L2/3 (~ 250 μm below brain surface) at delta-theta frequencies (2-8 Hz) exclusively in the Clemastine group. (B) The changes in Δcoherence between L4 and L2/3 over delta-theta frequency range was plotted over time, showing Clemastine administration resulted in significant elevation in L4-L2/3 connectivity compared to the vehicle control. (C) Heatmap of Δcoherence over depth paired with L5 electrode site as a function of frequencies. Clemastine administration resulted a strong coherence (white arrows) between L5 and L2/3 over alpha-beta frequency (8-30 Hz). (D) The changes in L2/3-L5 Δcoherence at alpha-beta band over time showed that Clemastine rescued the significant loss of L2/3-L5 connectivity at 14-16 weeks post-implantation. (E) L2/3-L5 mutual connectivity was quantified by Phase-amplitude coupling (PAC). Left: The feedforward projection from L2/3 to L5 was identified by the peak PAC modulation index between the phase of L2/3 slow theta band (4.5 - 7.5 Hz) and L5 55-70 Hz gamma amplitude (white arrow) in vehicle control mice. Right: the feedback projection from L5 to L2/3 was detected by the peak PAC modulation index between L5 theta phase and L2/3 60-70 Hz gamma amplitude (white arrow) in vehicle control mice. (F) The balance of L2/3 and L5 connectivity was quantified as the ratio of modulation index of L2/3 theta phase coupling with L5 gamma amplitude over that of L5 theta phase coupling with L2/3 gamma amplitude. Red arrows indicate significant differences between Clemastine-treated and vehicle control mice by a linear mixed-effects model following by likelihood ratio test with a 95% confidence interval.

Meanwhile, we also detected a strong functional connectivity between L5 and L2/3 exclusively in Clemastine-treated mice (Fig. 7C), indicated by the Δcoherence over alpha-beta frequency range (8-30 Hz). Clemastine group demonstrated a stable Δcoherence between L5 and L2/3 in alpha-beta band over time, whereas the vehicle group experienced a gradual reduction by nearly 134%. The resulting statistical difference (Fig. 7D, p < 0.05) between two groups indicates that Clemastine prevents the loss of functional connectivity between L2/3 processing layer to L5 information output layer over the chronic microelectrode implantation period.

Furthermore, there is a feedback loop between L2/3 and L5, with L5 neurons receiving inputs from L2/3 neurons and sending feedback up to L2/3 by direct anatomical synaptic connections [101, 102]. To further explore how Clemastine regulates the bidirectional connectivity between L2/3 and L5, we took advantage of the phase-amplitude coupling (PAC) analysis. The Hilbert transformation extracted the power and phase components of the LFP oscillatory data. Power represents the magnitude of neural activity at a specific frequency, and the phase provides the information related to the timing of neural activity. Integrating these two components, PAC describes the interactions between simultaneous LFP oscillations that occurred in different frequency bands, specifically how well the phase of an oscillation modulates the amplitude of another oscillatory signal at a different microelectrode channel (details in methods). Theta phase-gamma amplitude coupling has been reported as a hallmark of network functional connectivity during various cognitive tasks [103, 104], such as information processing and integration [105], and learning and memory formation [106].

Therefore, to characterize bidirectional functional connectivity between L2/3 and L5, we first calculated the PAC modulation index (MI) in vehicle mice to map the interlaminar theta phase-gamma amplitude coupling relationship bidirectionally. The value of MI was between 0 and 1, a larger MI value representing a stronger coupling between LFP low frequency phase and high frequency amplitude. Fig. 7E confirmed there was a feedback loop between L2/3 and L5 during visual network activation: L2/3 theta phase had a prominent coupling (modulation index: 1.398e-04 ± 7.153e-06) with 55-70 Hz amplitude at 600-750 μm depth corresponding to L5; in reverse, L5 theta phase was strongly coupled with 60-70 Hz amplitude in a modulation index of 1.913e-04 ± 7.264e-06 at L2/3 depth.

Then, we investigated whether the balance of L2/3-L5 feedback loop was disrupted by chronic microelectrode implantation and whether Clemastine treatment rescued the deficits of interlaminar connectivity. Modulation index ratio was used to quantify the balance of L2/3-L5 bidirectional connectivity, calculated as the modulation index of L2/3 theta-L5 gamma divided by the reverse direction L5 theta-L2/3 gamma. Fig. 7F showed that during initial 2 weeks postimplantation both Clemastine and vehicle group maintained the modulation index ratio close to 1, suggesting a balanced L2/3-L5 functional connectivity. Then, the vehicle group experienced a chronic decline in modulation index ratio since week 12 post-implantation, due to a reduced L2/3 to L5 feedforward connectivity or increased L5 to L2/3 feedback connectivity near the chronic implanted microelectrode. However, Clemastine mice showed a trend of the modulation index ratio near to 1 relative to vehicle controls (Fig. 7F). A significant difference (likelihood ratio test with non-overlapping 95% confidence intervals) in modulation index ratio between Clemastine and vehicle group was detected at week 15 postimplantation, indicating Clemastine helps restore the balance of L2/3-L5 bidirectional connectivity during the chronic microelectrode implantation. In summary, microelectrode implantation impairs functional interlaminar connectivity over the chronic implantation period, and promyelinating Clemastine treatment can rescue this interlaminar connectivity deficit.

### 3.3 Clemastine promotes oligodendrogenesis, remyelination, neuronal health near the chronically implanted microelectrode

At the end of the 16-week implantation period, brains were harvested and processed for immunohistochemistry. Cellular and subcellar markers were labeled in Clemastine-treated and vehicle mice tissue. We use this post-mortem immunohistochemistry to analyze how Clemastine affects oligodendrocyte population and neuronal health near the chronically implanted microelectrodes. Having demonstrated that Clemastine improves chronic electrophysiological recording performance, we investigated the relationship of this beneficial effect of Clemastine on recording performance to oligodendrocyte lineage structures starting with myelin. The fluorescent intensity of MBP, a marker for myelin sheaths, showed that Clemastine resulted in a significantly higher intensity up to 120 μm away from the probe relative to vehicle mice (Fig. 8A left, p < 0.0001). Additionally, MBP+ myelin in Clemastine group had a significantly higher level compared to the vehicle control on the contralateral hemisphere (Fig. 8A right, p = 0.0352). This MBP profile indicates Clemastine administration increases myelin in both implant injury and no-implant conditions.

**Figure 8:**
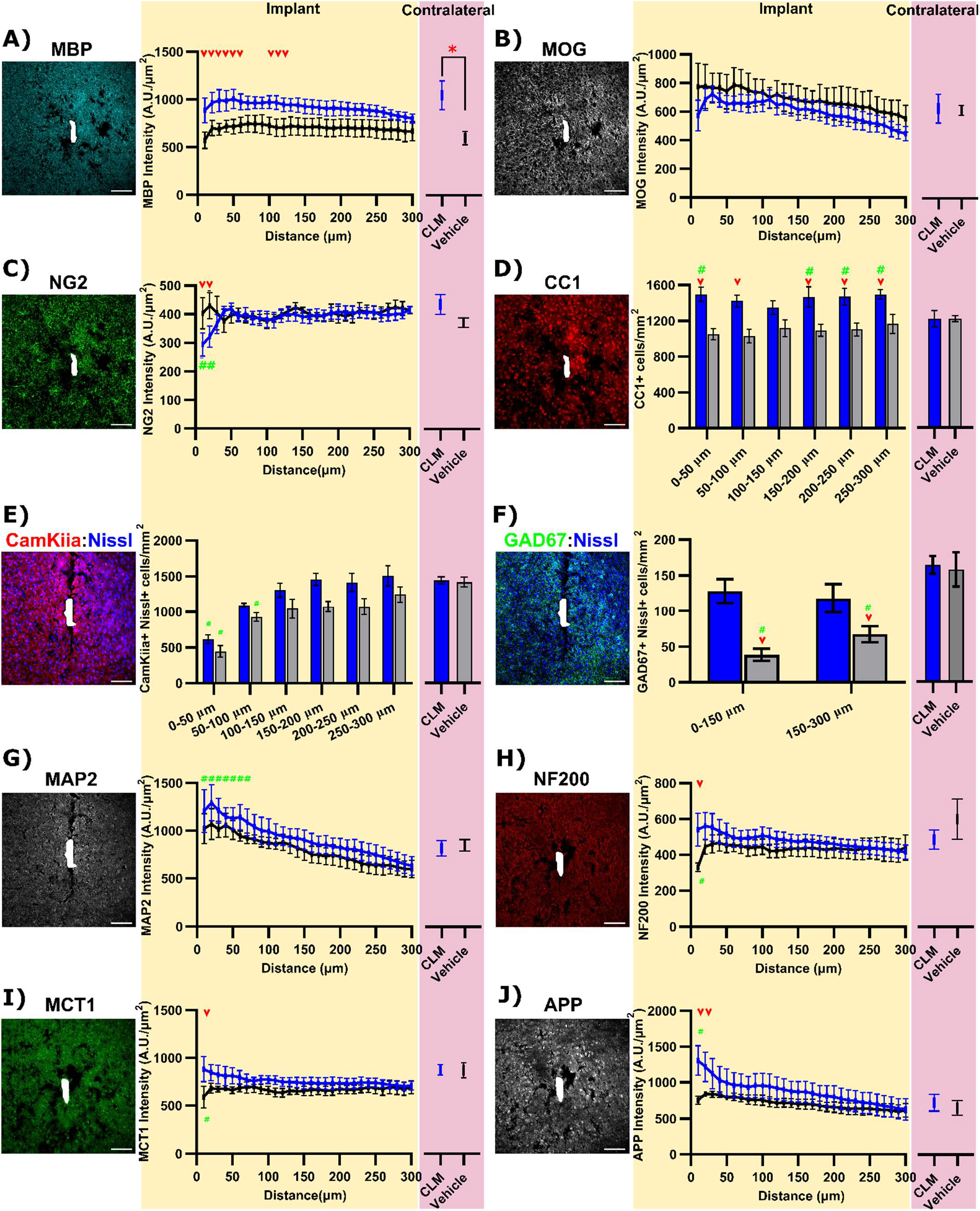
Clemastine daily administration increases oligodendrocyte activity and neuronal health by 16-week post-implantation. (A) MBP+ staining shows a global elevation in both implant site and contralateral region by 16-weeks Clemastine (CLM) administration. (B) MOG+ intensities are comparable between Clemastine mice and vehicle controls in both implant interface and contralateral regions. (C) Increases in CC1+ cells exclusively near the implant interfaces following Clemastine daily administration. (D) Intensity of NG2 fluorescence exhibits reductions up to 20 μm away from the probe in Clemastine group. Representative post-mortem tissue staining and cell count analysis of CamKiia+ Nissl+ excitatory neurons (E) and GAD67+ Nissl+ inhibitory neurons (F) between Clemastine (blue) and vehicle (black) mice. (G) Slight elevation in MAP2+ staining at implant site by 16-weeks Clemastine administration. Clemastine administration resulted increases in MCT1 (H), NF200 (I), APP(J) intensities at implant side compared to vehicle controls, while the expression in contralateral regions were comparable between Clemastine and vehicle mice. Red arrow indicates the group-wise significant differences in the implant side between Clemastine and vehicle conditions by two-way ANOVA and Fisher’s Least Significant Difference (LSD) post hoc (p < 0.05). Green # indicates the significant difference in fluorescent intensity between bins in the implant side compared to its contralateral. Red asterisks indicate significant different in contralateral fluorescence intensity between Clemastine and vehicle (unequalvariance Welch’s t-test). Scale bar, 100 μm.

Furthermore, the staining of an alternative myelin protein, myelin oligodendrocyte glycoprotein (MOG), provide details of enhanced myelination by Clemastine. MOG is located on the external lamellae of myelin sheath and associated with oligodendrocyte maturation and therefore, is usually present in relatively lower concentrations compared to other myelin proteins [107]. There were no significant differences in MOG+ intensities between Clemastine treated and vehicle controls in either bins at implant side (p > 0.05) or contralateral MOG intensity (p > 0.05) in Fig. 8B. Additionally, Clemastine mice demonstrated a slightly lower level in MOG+ fluorescence intensities relative to vehicle controls on the implant side. The distinct profile of MOG and MBP fluorescent intensity may depend on the properties of these two different myelin proteins. MBP acts as a major myelin protein and accounts for nearly 30% of the entire myelin proteins [108]. However, MOG is located at the outermost surface of multi-layer sheathed myelin, and therefore only accounts for a minor component (0.05%) of myelin composition [109]. In this way, while MBP can be used to examine the spatial distribution of myelin sheaths as well as myelin thickness, MOG is commonly used to evaluate the amount of mature myelin sheaths. The distinct MOG and MBP patterns imply that Clemastine administration may not affect the number of myelin segments but indeed increases the thickness of myelin sheaths. Alternatively, the slight reduction in Clemastine MOG intensities near the implant side suggests that there are more proportion of actively-growing myelin sheaths that are not mature yet, since the MOG levels are comparable in contralateral sides between two conditions.

Clemastine have been shown to promote oligodendrocyte precursor cells differentiation into mature oligodendrocytes [62]. We observed the expression of oligodendrocyte precursor cell marker, NG2, was significantly lower within the first 20 μm away from the probe in Clemastine group compared to vehicle controls (p < 0.005) as well as Clemastine no-implant contralateral (p < 0.005) in Fig. 8C. The CC1 marker for mature oligodendrocytes (Fig. 8D) demonstrated that Clemastine group exhibited significant increases in CC1+ cell density at the implant interfaces relative to vehicle controls (p < 0.0001). However, there was no significant difference in contralateral CC1+ cell density between Clemastine and vehicle regions (p = 0.9923). Meanwhile, comparisons in CC1+ oligodendrocyte density showed that there were significant increases in Clemastine near the implant relative to its contralateral side (p < 0.05), indicating that implantation injury resulted in the elevation in the number of mature oligodendrocytes. Overall, Clemastine administration effectively increases oligodendrocyte population and myelination at chronic 16-week microelectrode implantation time point.

Next, we focused on how Clemastine administration influences neuron density, which could support electrophysiological results. We examined the functional neuronal subtypes by colocalization of Nissl+ neuron nuclei with an excitatory soma marker, CamKiiα, and an inhibitory interneuron marker, GAD67. The density of CamKiiα+ Nissl+ cells was significantly reduced at 0-50 and 50-100 μm bins from the probe in the vehicle groups compared to contralateral regions (Fig. 8E, p < 0.05). In contrast, CamKiiα+ Nissl+ cell density was reduced only at the 0-50 μm bin in the Clemastine group compared to its contralateral hemisphere (Fig. 8E, p < 0.05), suggesting increased survival of excitatory neuron population near the implant. GAD67+ Nissl+ cells were significantly reduced in vehicle control mice approximately 76 % within 150 μm and 57% at 150-300 μm from the implant relative to its contralateral side (Fig. 8F, p < 0.0005). The number of GAD+ Nissl+ cells near the implant was significantly higher in the Clemastine treated mice compared to vehicle controls (p < 0.005) and was similar to Clemastine contralateral side (Fig. 8F, p = 0.7959). For neuron structural compartments, MAP2+ a marker for dendrites showed a slight fluorescence intensity increase in Clemastine mice compared to vehicle controls (Fig. 8G). The Clemastine mice showed a significant higher MAP+ intensity up to 70 μm compared to its contralateral, whereas the vehicle control had no significant difference between distance bins in implant side. Interestingly, the NF200+ axons in Clemastine group demonstrated a stable fluorescence distribution over distances from the implant. In contrast, the vehicle group showed a significant reduction in NF200+ intensity within 10 μm from the probe compared to Clemastine group (Fig. 8H, p < 0.0001). These MAP2 and NF200+ intensity profiles indicate that Clemastine helps preserve the neurite structural integrity during the chronic microelectrode implantation period.

Having shown that Clemastine increases OL and myelin density, we explored potential mechanisms of Clemastine mediated neuroprotection and measured metabolic components engaged in oligodendrocyte/myelin-neuron interactions. Metabolites delivery from myelin to axons is critical for axonal integrity [44]. Here, we assessed whether Clemastine affects expression of monocarboxylate transporter (MCT1) that mediates the delivery of metabolites (e.g. lactate and pyruvate) from myelin to axons. Clemastine administration resulted in a significant elevation in MCT1 expression within 10 μm away from the implant relative to vehicle controls (Fig. 8I, p < 0.005). The signal intensity of amyloid precursor protein (APP) that plays a critical role in synaptic plasticity [110] showed a significant increase near the implant in Clemastine group compared to vehicle mice, specifically within 20 μm from the probe (Fig. 8J, p < 0.0001). In summary, Clemastine that promotes oligodendrogenesis near the implant has neuroprotection effect, preserving the soma of excitatory and inhibitory neurons, increasing axonal integrity, and reducing metabolic deficits resulted by implantation injury relative to vehicle controls.

**Supplementary Figure 5:**
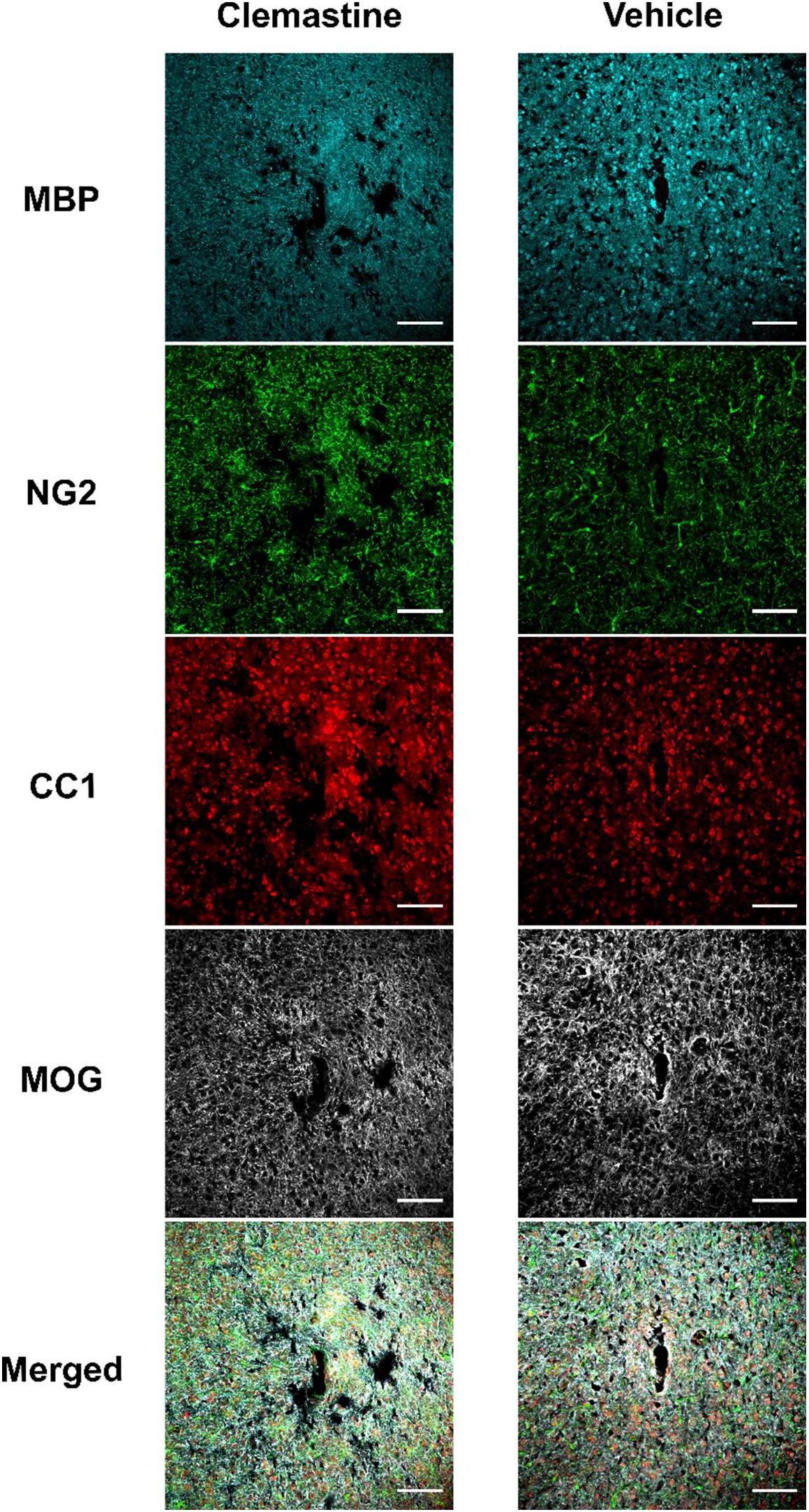
Representative histological images of oligodendrocyte lineage structures in Clemastine and vehicle tissue. Staining combination of MBP, NG2, CC1, MOG and an overlayed image to investigate oligodendrocyte and myelin activity between Clemastine and vehicle tissue. Scale bar, 100 μm

**Supplementary Figure 6:**
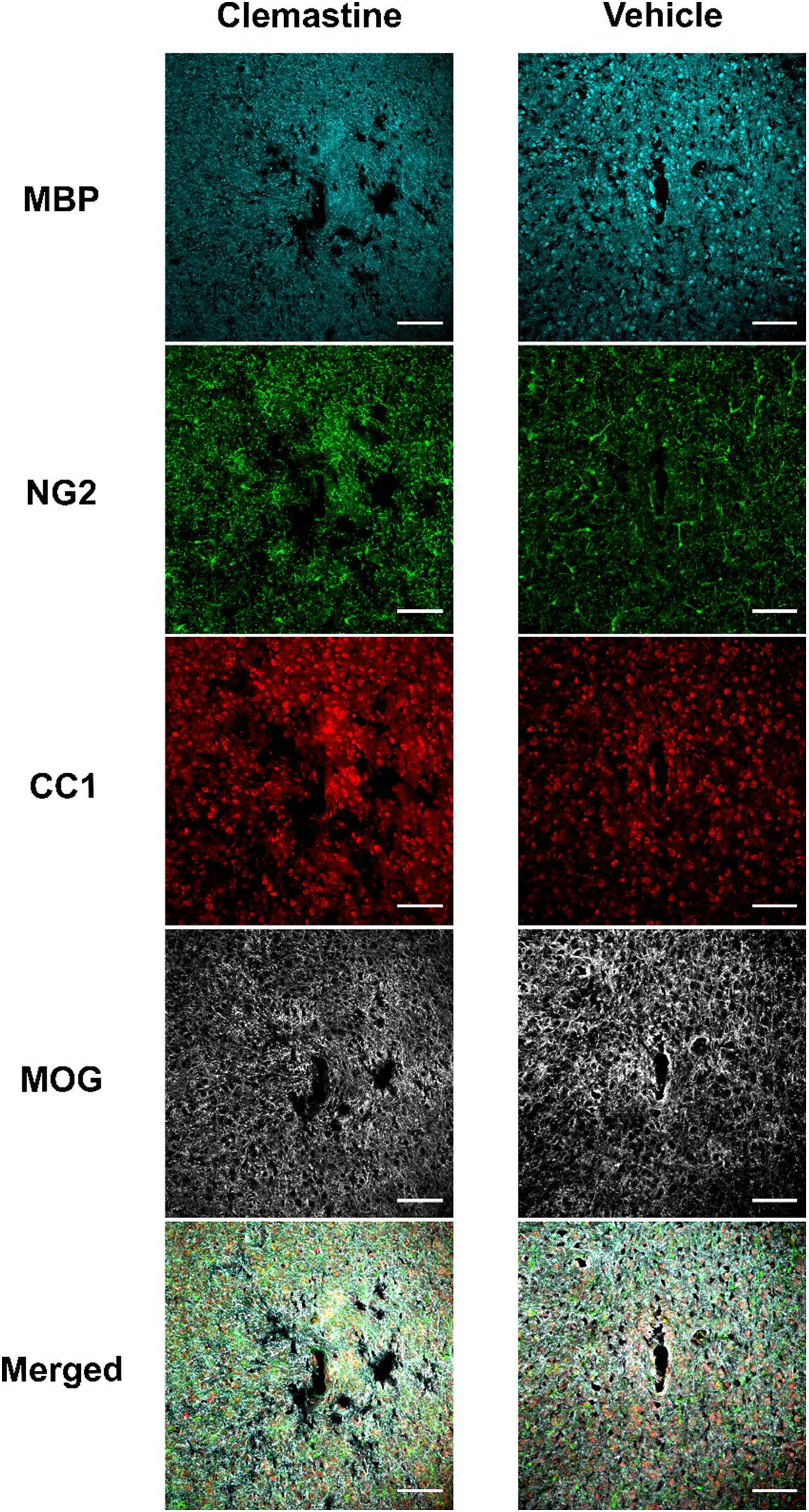
Representative histological images of neuron subtype distributions in Clemastine and vehicle tissue. Staining combination of Nissl, GAD67, CamKiia, MAP2, and an overlayed image to understand whether neuronal subpopulation were changed between Clemastine and vehicle mice. Scale bar, 100 μm.

**Supplementary Figure 7:**
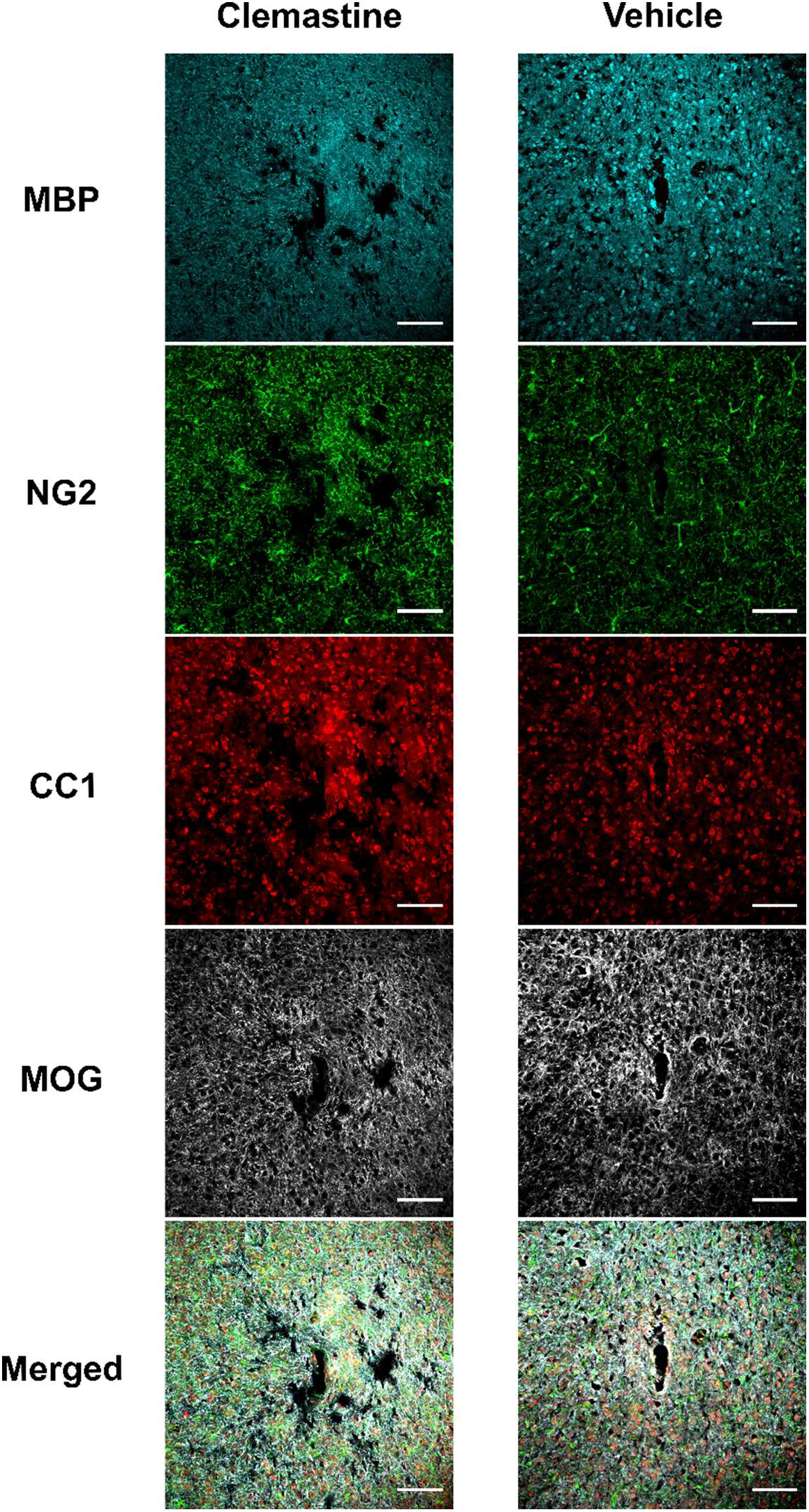
Representative histological images of subcellular markers in Clemastine and vehicle tissue. Staining combination of MBP, MCT1, NF200, APP and an overlayed image to understand the metabolic and plasticity mechanism by Clemastine administration. Scale bar, 100 μm.

## 4. Discussion

The impact of enhancing oligodendrocyte activity for improving long-term electrophysiological recording stability and sensitivity was investigated by using a pro-myelination therapeutic drug, Clemastine. Clemastine selectively affects oligodendrocyte lineage cells, promoting their differentiation into mature myelinating oligodendrocytes [62]. Additionally, side-effects from Clemastine are low, rarely including mild drowsiness fatigue, and dizziness, which fade with continued use as tolerance develops [82]. Single-shank Michigan-style microelectrode arrays were chronically implanted to record neural activity in visual cortex and hippocampus over 16 weeks in Clemastine and vehicle treated mice. Electrophysiological activity and histology revealed a positive correlation between enhanced functional recording performance and oligodendrocyte activity over the chronic implantation period. Specifically, Clemastine administration resulted in increased chronic recording performance, including increasing SU detection and firing rate, preventing the chronic loss of MU firing rates, improving the interlaminar LFP oscillatory connectivity. Furthermore, Clemastine helps increase the survival of excitatory and inhibitory neurons near the implant as well as rescue the damage of neuronal compartments. These results indicate that the beneficial effects of Clemastine on chronic recording performance arise from the increased neuronal integrity near the microelectrodes. Therefore, our findings suggest that therapeutic strategies enhancing oligodendrocyte and myelin activity could enhance the integration of brain tissue to implanted microelectrodes and mitigate the failure of functional devices.

### 4.1. Oligodendrocyte contribution to energy metabolism

The recording performance fidelity of chronically implanted microelectrodes often dictate the performance limits of prosthetic systems in clinical applications and of interrogating discrete brain activity in neuroscience studies [111]. Here, we demonstrated that promyelination pharmacological treatment mitigated the loss of signal quality in SU activity (Fig.1–2), MU firing rate (Fig. 3), and LFP oscillation power (Fig. 4). We showed that the increased recording performance was matched with increased oligodendrocyte population (Fig. 8C) and myelination (Fig. 8A-B) by Clemastine, which resulted in enhanced neuronal survival near the microelectrode (Fig. 8E-F). Enhanced oligodendrocyte activity has implications on increasing neuronal metabolic supply, which is highlighted by growing evidence regarding the role of myelination on supporting axonal metabolism [42]. This is supported by the enhanced MCT1 profile (Fig. 8I) indicating an increased metabolite transportation between myelin and axons near the chronic implanted microelectrode. In turn, this reduces the impairment of firing activity during increases in metabolic demand related to the progression of the foreign body response. Taken together, Clemastine may improve chronic recording performance by supporting oligodendrocytes and myelin to increase metabolic supply to neurons to protect their health and functionality.

However, it is important to note that MCT1 is not exclusively expressed in oligodendrocytes and myelin, but is also expressed in astrocytes as well [112, 113]. Astrocytes can shuttle lactate to neurons at the nodes of Ranvier [114], or to oligodendrocytes through gap junctions [115, 116]. The disruption of gap junctions connecting oligodendrocytes and astrocytes has been shown to diminish the axonal firing [117], which highlight the importance of these gap junction in metabolite trafficking between oligodendrocyte to astrocytes. Following microelectrode implantation, astrocyte process migration occurs toward the probe within the first 7 days [118]. This activation may disrupt the oligodendrocyte-astrocyte gap junctions and result in fluctuation of firing activity (Fig. 1–4). Then, at chronic 2-4 weeks, astrocytes undergo hypertrophy and ultimately form an encapsulating glial scar [118], while oligodendrocytes and myelin progressively degenerate of near the microelectrode [59]. A disruption of oligodendrocyte-astrocyte coordination would lead to the dysfunction of metabolic support to neurons over the chronic implantation period, which could explain the overall loss of MCT1 (Fig. 8I) and gradual loss of recording performance (Fig. 1–4) in vehicle mice. However, it is still unknown how chronic implantation injury disrupts oligodendrocyte-astrocyte coordination as well as the exact metabolic contribution of oligodendrocytes and astrocytes on axonal health. Future studies should focus on how Clemastine alters this cellular metabolic support.

Alternatively, the overall increase in MCT1 expression caused by Clemastine (Fig. 8I) could imply increased metabolic demand by enhanced oligodendrocyte activity. Neurons are not the only metabolically expensive cell type in the brain [119]. Oligodendrocytes also require considerable energy or metabolic costs to differentiate from OPC as well as produce and maintain myelin sheaths [117]. However, the inflammatory environment caused by chronic microelectrode implantation may lead to metabolic stress and local nutrient deprivation, due to the reduced metabolic supply such as loss of perfusion [35] as well as increased metabolic consumption by microglia activation and astrocyte reactivity [120, 121]. It is likely that Clemastine administration shifts the energy utilization to oligodendrocyte differentiation and myelination, leading to increased number of mature oligodendrocytes and myelin density(Fig. 8A-C). The decreased intensity of OPC marker NG2 in Clemastine mice (Fig. 8D) likely indicates OPC differentiation where NG2 expression is gradually diminished [60]. Additionally, enhanced myelination by Clemastine may increase the efficiency of energy utilization and metabolic costs of underlying axons. Increased myelination substantially reduce the amount of metabolic costs that a neuron requires following an action potential. Saltatory conduction by myelination limits ion flow across the extracellular membrane during depolarization to the nodes of Ranvier and therefore requires less energy to pump out ions and repolarize the membrane after each action potential along the axon [122]. Therefore, the Clemastine induced increase in metabolic cost to increase oligodendrocyte activity maybe outweighed by the metabolic savings from repolarization during a period of increased metabolic deficit due to increased metabolic demand from neuroinflammation and decreased metabolic supply form blood-brain barrier injury. This implication is further supported by our results of enhanced neuronal firing activity (Fig. 5) and increased neuronal survival (Fig. 8E-F) in Clemastine mice over the neuroinflammatory and chronic foreign body response period.

Furthermore, increased oligodendrocyte activity may help neurons survive against neurodegeneration. Following the microelectrode implantation, there is an increased gradient of reactive oxygen species, proinflammatory cytokines, and excessive glutamate release near the microelectrode [16, 123]. In this neuroinflammatory environment, unmyelinated axons likely become vulnerable to injury or degeneration. However, the presence of myelin helps restrict the diffusion of pro-inflammatory molecules at the paranodal junctions [44], reduce the exposure of axons to proinflammatory insults, and continue to provide neurotrophic factors [124]. The significant elevation of NF200+ intensity in Clemastine mice relative to control (Fig. 8H) indicates preserved axonal integrity near the microelectrode by increased oligodendrocyte activity. Moreover, the metabolites released from oligodendrocyte are stimulated by activation of glutamate NMDA receptors [125], and may facilitate metabolic support for neuronal firing activity. Also, the elevated SU yield (Fig. 1C) and signal amplitude (Fig. 1F) in Clemastine mice further suggest that the signal transduction along axons experience less damage due to increased myelination. This effect may have been enhanced by the 7-day treatment by increasing OPC differentiation and myelination prior to the implantation injury, although this needs to be further investigated in the future. Taken together, our results show that Clemastine provide therapeutic benefits to chronic recording quality and add support to the literature that oligodendrocytes contribute to neuronal integrity and functionality by providing metabolic support to nearby neurons.

### 4.2. Oligodendrocyte integrity mitigate damage in lipid metabolism

The brain is the second most lipid-rich organ, so the lipid metabolism is tightly related to development and maintenance of brain health and function [126]. However, excessive accumulation of lipotoxic metabolites by impaired lipid metabolism is linked to various neurological disorders such as multiple sclerosis (MS) [127], brain ischemia [128], and amyotrophic lateral sclerosis [129]. Oligodendrocytes generate lipid-rich myelin sheaths, which accounts for ~ 40% of the total lipids synthesized in the human brain [128, 130]. For example, cholesterol is a major myelin lipid that is required for myelin compaction [130]. Chronic implantation leads to progressive damage to myelin sheaths [59], which likely leads to abnormally high level of lipids and exacerbates local oxidative stress, inflammation, and ultimately neuron death [131]. However, Clemastine helped preserve myelin integrity (Fig. 8A-B) and demonstrated a positive therapeutic effect on electrophysiological measurements (Fig. 1–7). These observations suggest a novel perspective that improving the integrity of lipid metabolism near the implant may help contribute to neuronal health and functionality.

Clemastine administration elevated myelination as indicated by a significant increase in MBP (Fig. 8A) and a similar increase in MOG (Fig. 8B) relative to vehicle controls. Previous evidence showed that implantation injury upregulated Apolipoprotein E (ApoE) throughout the chronic 6-week implantation period [72], which has been associated with the transport of cholesterol and other lipids [72]. Since myelin sheaths are progressively degraded during chronic implantation injury [59], this loss of myelin integrity may lead to spread of lipid-containing myelin debris in the local tissue near the implant. The long-term accumulation of lipid (lipotoxicity) is deleterious to neuronal health, including destabilization the membrane integrity, damage to mitochondria respiration, production of reactive oxygen species [132]. These inflammatory processes are similar to neuroinflammatory response caused by microelectrode implantation [123]. The irreversible decline in recording performance (Fig. 1) in vehicle control animals highlight the dysfunction of neuronal activity, which could be the result of lipotoxicity surrounding the microelectrode. However, Clemastine treatment protects against myelin degradation and thus may reduce the amount of free lipids in the environment near the implant, which, in turn, mitigates the lipotoxicity-induced neuroinflammation. The improvement in neuronal health is demonstrated by the significant elevation in neuronal electrophysiological activity of Clemastine animals relative to vehicle controls (Fig. 1–7).

Additionally, the positive therapeutic effect of Clemastine could be benefits from appropriate lipid activity during myelination. Lipid is a required component for a mature myelinating oligodendrocytes [133]. Disruption in lipid metabolism result the improper lipid composition of myelinating oligodendrocyte, which leads to failure of oligodendrogenesis from OPC [133]. Previous studies showed oligodendrocyte degeneration outpaced oligodendrogenesis leading to remyelination failure near chronically implanted microelectrodes [59]. However, Clemastine has been shown to inhibit enzyme emopamil-binding protein (EBP) and ultimately increase cholesterol synthesis [134]. It is worth mentioning a potential alternative mechanism of Clemastine which may be increased OPC differentiation into myelinating oligodendrocytes. The myelin profile demonstrates a significant increase in MBP but a comparable level in MOG (Fig. 8B) in Clemastine mice relative to vehicle controls, which suggests Clemastine likely increase the thickness of myelin sheath rather than the amount of myelin segments. This potential increase in sheath thickness may rely on involvement of cholesterol in myelin multi-layer compaction. Moreover, our results of the promoted neuronal firing activity (Fig. 1–7) further indicate that successful myelination is functionally integrated with axons. Thus, it may be worth understanding whether synthesis of cholesterols in myelin is emerging as a key factor of neuroprotection during inflammatory environment. Future studies would help to develop novel therapeutic strategies that promote remyelination through modulating oligodendrocyte lipid metabolism.

### 4.3 Oligodendrocyte regulate the balance of excitatory and inhibitory network

Our results indicate that enhanced oligodendrocyte activity leads to promote excitatory-inhibitory network balance in activated visual circuit. Increased survival of excitatory neuron density (Fig. 8E) as well as concomitant sustained firing rate (Fig. 5E) suggests that the excitatory network is improved with the promyelinating Clemastine administration compared to vehicle controls. While there is substantial myelin coverage over excitatory axons, some oligodendrocyte also myelinate inhibitory PV axons [135]. It has been proposed that myelin regulates PV+ neurons metabolism as well as improves the energy efficiency of signal propagation [136–138]. Thus, the neuroprotective effect of promyelinating Clemastine were also observed in inhibitory networks near the implanted microelectrode. In Clemastine treated mice, the observed firing activity of putative inhibitory SU (Fig. 5I-J) correlated with an enhanced density of GAD+ inhibitory neurons (Fig. 8F) indicating the preservation of inhibitory neuron network. Thus, our results suggest a tight correlation between the level of oligodendrocyte activity, myelination, and neuroprotection of excitatory/inhibitory networks.

The instability of chronic recording signal quality can be attributed to the shift in the balance of excitatory/inhibitory tone at the microelectrode interfaces over time. Previous study revealed a gradual loss of excitatory VGLUT1 marker with a concomitant increase in inhibitory VGAT marker over a 4-week microelectrode implantation period [139]. This loss of excitatory tone is similarly indicated by the reduced power of slow oscillations that likely reflect synchronous excitatory synaptic activity. The promyelinating Clemastine significantly rescued this power loss over delta (Fig. 4E) and theta bands (Fig. 4F), indicating that the excitatory network is preserved by enhanced myelination. Additionally, oligodendrocytes can modulate inhibitory network with collaboration with astrocytes. The ability of astrocytes to amplify GABAergic inhibition of pyramidal neurons [140] may be influenced by the oligodendrocyte activity due to the tight oligodendrocyte-astrocyte coupling [141]. Thus, the enhanced oligodendrocyte activity by Clemastine likely contributes to inhibitory network functionality in coordination with astrocytes. However, the role of astrocyte in Clemastine’s neuroprotection is not investigated in this study. Future works may focus on the contribution of additional glia participate in the regulation of excitatory/inhibitory network surrounding the microelectrode.

Furthermore, synapse plasticity is also critical to excitatory/inhibitory network balance, which has been closely related to the level of Amyloid Precursor Protein (APP). Thus, the APP level near the microelectrode in Clemastine and vehicle mice (Fig. 8J) reflects the level of synapsic connections in the functional excitatory/inhibitory network. Although APP upregulation is known in pathogenesis of Alzheimer’s disease (AD), recently emerging evidence emphasize the role of this protein in synaptic transmission, plasticity, dendritic sprouting, and calcium homeostasis [110, 142] for normal function. APP regulates the postsynaptic glutamatergic signaling [143] as well as presynaptic GABA receptor functions [144]. As the increases in APP improve the expression and function of N-methyl-D-aspartate receptors (NMDAR) receptors [143], the elevated APP intensities near the implant by promyelination Clemastine (Fig. 8J) indicate enhanced excitatory network relative to vehicle condition. Also, APP is highly expressed in GABAergic interneurons and regulates phasic and tonic inhibition [145]. Therefore, this increase in APP intensity profile at Clemastine implant site (Fig. 8J) may suggest a less-influenced inhibition that could be balanced well with excitation. Additionally, APP can facilitate synaptic plasticity by modulating glutamatergic NMDAR and multiple calcium channels that are critical for long term potentiation involved in learning and memory [110]. A previous study has shown that long-term implantation over motor cortex induces onset of behavioral deficits [146], which implies a failure of synaptic activity in the network. Taken together, the enhanced APP expression by promyelinating Clemastine near the chronic implant site (Fig. 8J) is likely to mitigate the impairment of synaptic connectivity, which is further supported by the increased electrophysiologically detected interlaminar connectivity (Fig. 7).

### 4.4 The region-specific effect of Clemastine’s neuroprotection

Interestingly, we observed that the level of neuroprotection from Clemastine is dependent on brain regions. The cortex maintained elevated recording quality throughout the 16-week implantation period, whereas in the CA1 region, performance declined to control level by 10 weeks post-implantation (Fig.2). This distinction implies that Clemastine has a different impact on neuronal health and functionality in different brain circuits. The heterogeneity of oligodendrocyte in morphology, gene expression, and myelination activity in cortex and hippocampus has been reported, which highlight their diverse roles in functional neural activity of different brain region [147–149]. Oligodendrocytes in hippocampus have been suggested to have a longer timing for maturation and myelination compared to those in the cortex during development [150, 151]. Thus, Clemastine may result in a slower progress in oligodendrocyte differentiation in hippocampus and thus have a relative mild neuroprotection compared to cortex. However, the tissue injury in hippocampus is likely more severe than cortex due to more mechanical strain located at deeper depth [8]. Thus, the mild positive effect by Clemastine may eventually be overwhelmed by the neuroinflammatory and foreign body response in the hippocampus, which can lead to the chronic decline in CA1 recording quality of Clemastine mice. Future studies could focus on understanding the influences of different oligodendrocyte subpopulations on neural activity, which help further reveal the mechanism of oligodendrocyte – neuron functional coupling.

The different vasculature in cortex and hippocampus is another possible factor of region-specific impact by Clemastine. Supply of glucose and oxygen by cerebral microcirculatory blood flow is tightly associated with the level of metabolism in local tissue [152]. Chronic administration of Clemastine drives additional oligodendrocyte differentiation and myelination (Fig. 8), which require a large and continuous energy supply. However, the microvascular networks, which are the primary source of metabolic supply in the brain, are distinct between cortex and hippocampus [153]. The neurovasculature in CA1 hippocampus has a lower mean vascular diameter, volume fraction, and length density compared to cortex [153, 154]. Moreover, the vessel architecture in hippocampus farther away from the surface large arterioles and has a higher blood flow resistance [155], which increase the difficulty of metabolic delivery to CA1 relative to cortex. Thus, the hippocampus is more sensitive to experiencing vulnerability to small metabolic deficits to persistent implantation injury compared to cortex. Therefore, it is possible that Clemastine is insufficient in promoting oligodendrocyte differentiation and myelination in CA1 hippocampus under environments stressed by neuroinflammatory and foreign body responses. The chronic recording failure in CA1 despite Clemastine treatment suggest that alternative pathways may need to be explored for improving recording performance in the hippocampus.

Alternatively, it is possible that microelectrode implantation results in different levels of impairment in information input for circuit activation in cortex and hippocampus, respectively. Visual cortex is able to receive driving input vertically from subcortical regions [156] such as thalamus as well as horizontally from other cortical regions [157]. Although the microelectrode was perpendicularly penetrated into the brain, cortical circuits near the implant may still be strongly activated through the horizontally connecting axons from adjacent regions. Instead, CA1 hippocampus receives cortical inputs through the cingulum fiber bundle, a special white matter tracts in a ring shape that linking multiple cortical sites to subcortical regions [158]. Dysfunction in cingulum bundle is associated with hippocampus atrophy in neurodegeneration disease [159]. The perpendicular implantation of microelectrode arrays into CA1 also damage the cingulum bundle above the CA1. This injury to the overlying cingulum bundle likely impairs the sensory information relayed from visual cortex to hippocampus. This impairment in hippocampus afferents may contribute to the continuously declining CA1 recording performance, and the chronic damage to the cingulum bundles could overwhelm the Clemastine’s neuroprotective effects for recording CA1 activity.

### 4.5 Limitations and future directions

A few limitations exist in the study due to the restrictions of experimental designs and inherent data collection procedures. First, we do not know if the one-week Clemastine preconditioning prior to microelectrode implantation is necessary. Our results demonstrated the daily Clemastine with one-week preconditioning leaded to significant difference in SU detection since day 6 post-implantation. However, to what extent of neuroprotection effect that Clemastine preconditioning may have was not specifically explored in this study.

The second limitation is the unknown contribution of pre-existing oligodendrocyte and newly differentiated oligodendrocyte to enhanced myelination by Clemastine. Previous study has shown the inflammatory environment caused by the microelectrode implantation impairs the oligodendrocyte differentiation activity [59]. Here, it is unclear that whether the ability of pre-existing and newly differentiated oligodendrocyte to produce myelin are impacted by 16 weeks chronic implantation. However, regardless of these underlying pathways Clemastine demonstrates an enhanced oligodendrocyte activity near the microelectrode and results a robust chronic recording performance compared to vehicle controls.

Another major limitation is that how other glial cells are influenced by Clemastine near the chronic implant is not known. While this study focuses on the influences of oligodendrocyte activity on neuronal integrity over chronic Clemastine administration, microglia and astrocytes cross-talks with oligodendrocytes [141, 160, 161]. The microglia dynamics tightly regulates myelination phases, including degeneration and remyelination, during pathological conditions [160, 162]. In lactate shuttle theory, astrocytes that have gap junctions with oligodendrocytes facilitate metabolic support of oligodendrocytes to neurons [141]. The influences of Clemastine administration on microglia activation and astrocyte reactivity to the chronically implanted microelectrodes remain unknown. However, the emerging evidence shows that the relationship among microglia, astrocytes and oligodendrocytes is complex [161], which suggests that a separate study is required for detailed characterizations of the impact of enhanced oligodendrocyte activity on microglia activation and astrocyte reactivity. Therefore, researchers interested on glial activity can focus on the interactions of oligodendrocyte, microglia, and astrocyte activity at brain-machine interfaces over time.

One future direction is to provide spatiotemporally mapping of Clemastine’s influences on the chronically implanted microelectrode. Here, the histological staining results only provide the one time point snapshot of tissue changes at the end of 16 weeks implantation. The limited temporal characterization of tissue changes by Clemastine impedes the understanding the role of oligodendrocytes in functional integration of the implanted microelectrodes into the brain circuit. Therefore, alternative methods such as two-photon in vivo microscopy can be used to explore when, where, and how Clemastine affects oligodendrocyte activity between degeneration and regeneration. Specifically, transgenic models that can simultaneously express oligodendrocytes and neurons can further reveal the dynamics of oligodendrocyte-neuron coupling during implantation-induced injury under Clemastine administration.

Another future direction could be the mechanistic investigation of Clemastine on oligodendrocyte-neuron coupling near the chronic implant. Clemastine has been reported to affect oligodendrocyte lineage structure without significantly changing neuronal/axonal density [62]. However, our results showed enhanced oligodendrocyte lineage activity and improved excitatory and inhibitory neuronal density at the end of 16 weeks post-implantation. These differences may reflect the ability of Clemastine to rescue neurons in greater metabolic stress environments such as near chronically implanted microelectrodes compared to hypoxia only. Although the MCT1 and APP staining provides some potential perspectives, specific knockout or mutation models such as oligodendrocyte MCT1 knockout mice should be considered to disentangle the how myelination promoted by Clemastine protect the neuronal dysfunctions near the chronically implanted microelectrodes.

## 5. Conclusion

Enhanced oligodendrocyte activity by Clemastine administration promoted the recording performance of chronically implanted microelectrode and improved functional neural activity in the surrounding brain area. Clemastine administration resulted in elevated SU and multi-unit metrics following the initial 2-week inflammatory period and remained robust throughout the entire 16-week implantation period. Specifically, Clemastine increased the viability and firing properties of both excitatory and inhibitory neurons. Additionally, Clemastine rescued the degradation of global network connectivity during the chronic phase of implantation as well as prevented the loss of oscillatory activity and interlaminar functional connectivity. Overall, this study demonstrated the feasibility of a novel therapeutic strategies targeting oligodendrocyte activity for improving chronic recording performance. These findings reveal the importance of oligodendrocyte function on the functional integration of chronically implanted microelectrodes to the brain tissue.

## Acknowledgement

The authors would like to thank Steven Wellman, Kevin Stieger, Naofumi Suematsu, Fan Li, Camila Garcia for assistance with manuscript proofread.

